# Chemokine receptor 5 signaling in PFC mediates stress susceptibility in female mice

**DOI:** 10.1101/2023.08.18.553789

**Authors:** Hsiao-Yun Lin, Flurin Cathomas, Long Li, Romain Durand-de Cuttoli, Christopher Guevara, Cigdem Sevim Bayrak, Qian Wang, Swati Gupta, Kenny L. Chan, Yusuke Shimo, Lyonna F. Parise, Chongzhen Yuan, Antonio V. Aubry, Fiona Chen, Jean Wong, Carole Morel, George W. Huntley, Bin Zhang, Scott J. Russo, Jun Wang

## Abstract

Chronic stress induces changes in the periphery and the central nervous system (CNS) that contribute to neuropathology and behavioral abnormalities associated with psychiatric disorders. In this study, we examined the impact of peripheral and central inflammation during chronic social defeat stress (CSDS) in female mice. Compared to male mice, we found that female mice exhibited heightened peripheral inflammatory response and identified C-C motif chemokine ligand 5 (CCL5), as a stress-susceptibility marker in females. Blocking CCL5 signaling in the periphery promoted resilience to CSDS. In the brain, stress-susceptible mice displayed increased expression of C-C chemokine receptor 5 (CCR5), a receptor for CCL5, in microglia in the prefrontal cortex (PFC). This upregulation was associated with microglia morphological changes, their increased migration to the blood vessels, and enhanced phagocytosis of synaptic components and vascular material. These changes coincided with neurophysiological alterations and impaired blood-brain barrier (BBB) integrity. By blocking CCR5 signaling specifically in the PFC were able to prevent stress-induced physiological changes and rescue social avoidance behavior. Our findings are the first to demonstrate that stress-mediated dysregulation of the CCL5-CCR5 axis triggers excessive phagocytosis of synaptic materials and neurovascular components by microglia, resulting in disruptions in neurotransmission, reduced BBB integrity, and increased stress susceptibility. Our study provides new insights into the role of cortical microglia in female stress susceptibility and suggests that the CCL5-CCR5 axis may serve as a novel sex-specific therapeutic target for treating psychiatric disorders in females.

---

Stress disorders are leading causes of disability with significant health implications. Both genetic and environmental factors influence the risk of developing stress disorders across the lifespan. Psychosocial stressors have been shown to increase peripheral cytokine production, which may contribute to the development of depression and anxiety in humans^1,2^. In particular, subsets of patients with major depressive disorder (MDD) exhibit elevated levels of systemic inflammatory markers, including cytokines IL-6, TNF-α, CRP, and MCP-1^3–5^. Meta-analyses have also indicated that proinflammatory cytokines are associated with the progression and severity of depressive disorders across different populations^6^ and influence their responses to treatment^7^.

Neurovascular dysfunction and blood-brain barrier (BBB) disruption have also been linked to the pathogenesis of stress disorders^8–11^. Clinical evidence suggests that cerebrovascular disease may predispose select populations to stress disorders; for example, the prevalence of MDD is 2-3 times higher in patients with cardiovascular diseases and is correlated with increased morbidity and mortality^12–14^. Late-life depression in older adults is consistently associated with greater white matter lesions (indicative of vascular dysregulation) and increased expression of adhesion molecules (indicative of inflammation) in various brain regions including the dorsolateral prefrontal cortex (DL-PFC) and anterior cingulate cortex (ACC)^15–18^, suggesting potential interactions between peripheral inflammation, vascular dysfunction, and depression^19^.

Systemic inflammation and vascular dysfunction may lead to an increase in peripheral immune infiltration into the central nervous system (CNS), where they can directly interact with CNS-resident cells such as astrocytes, microglia, and neurons. Microglia, which constitute 10% of all brain cells, are the major type of immune cells in the CNS. They play important roles in normal development, maintain brain homeostasis by actively surveilling cells, eliminate synapses and other cellular debris, and promote neurogenesis^20^. Under pathological conditions, microglia can exhibit a broad spectrum of activation states in response to various stimuli, including morphological changes, production of inflammatory cytokines^21^ and increased phagocytic capacity. Notably, the number of activated microglia is significantly higher in subjects with various psychiatric disorders compared to healthy controls in postmortem brains^22–25^. Recent imaging studies have detected increased microglial activation in the PFC, insula, and ACC of human subjects with psychiatric conditions, including depression^26–29^, with microglial activation being positively correlated with the severity of depression^30^.

Preclinical stress studies largely recapitulated what is observed in human stress disorders, including heightened peripheral inflammation, vascular pathology, altered BBB permeability and neuronal remodeling^31–35^. Microglial activation following inescapable shock^36^, social defeat stress^37^, and chronic unpredictable stress has also been reported in mice^38^, and in restrained rats^39,40^. However, the interactions between the peripheral immune system, cerebral vasculature, and CNS resident cells under stress conditions are largely unknown. More importantly, the majority of preclinical studies have been conducted in males only. Sex differences in psychiatric disorders, including anxiety, depression, and addiction, have been well-documented, with women generally exhibiting higher vulnerability in terms of prevalence, symptom persistence and severity, and responses to treatment^41^. Therefore, knowledge gaps exist regarding female-specific immunological, electrophysiological, and neurobehavioral responses to stress exposure.

The current study investigates how peripheral inflammation induced by stress influences microglial activity in the PFC in a female chronic social defeat stress (CSDS) model, and the participation of activated microglia in excessive phagocytosis of neuronal and vascular material, leading to alterations in neurotransmission and BBB pathology, which may ultimately contribute to stress-related behavioral changes.

## Peripheral CCL5 levels correlate with stress-susceptibility in female mice

Dysregulation of immune response has been linked to stress disorders including depression. In previous studies in male mice, CSDS induced significant elevation of the pro-inflammatory cytokine IL-6 in the periphery and its level correlated with susceptible phenotype^42,43^. To study peripheral inflammatory responses to stress in female mice, we exposed female mice to 10 days of CSDS followed by social interaction (SI) behavior test. Blood was collected via the submandibular vein one day after the first defeat bout to assess for initial immune responses (**Fig. 1A**). Following 10-day CSDS and SI behavior testing, mice were categorized as stress-susceptible (SUS) or resilient (RES) based on the SI ratio, which is calculated as the time spent in the interaction zone with a target mouse divided by the time spent in the interaction zone without a target mouse. We found a significant decrease in SI ratio and increase in time spent in the corner zones in defeated mice (**Fig. 1B and 1C**). Peripheral cytokines one day after the first defeat bout were analyzed by multiplex Enzyme-linked immunosorbent assay (ELISA). Compared to cytokine profiles from male mice after the first defeat bout, we found female mice demonstrated heightened pro-inflammatory responses with significantly higher levels of CCL5, IL-1α, IL-1β, IL-6, IL-12(p70), TNF-α, and MCP-1 compared to non-defeated control mice while in male mice, only IL-6 was significantly increased (**Extended Data Table 1**). This is consistent with literature suggesting that there are sex differences in the immune system that might make females more vulnerable to inflammatory diseases^44,45^. Correlation analysis between plasma cytokine/chemokine levels and SI/corner zone occupancy showed that unlike male mice where SI ratio negatively correlates with plasma levels of IL-6^46^, in female mice, SI ratio was negatively correlated with plasma levels of CCL5 (**Fig. 1D**) while corner zone occupancy duration was positively correlated with CCL5 (**Extended Data** Fig. 1A). None of the other differentially regulated cytokines showed correlation with SI and corner zone occupancy.

**Fig. 1.**
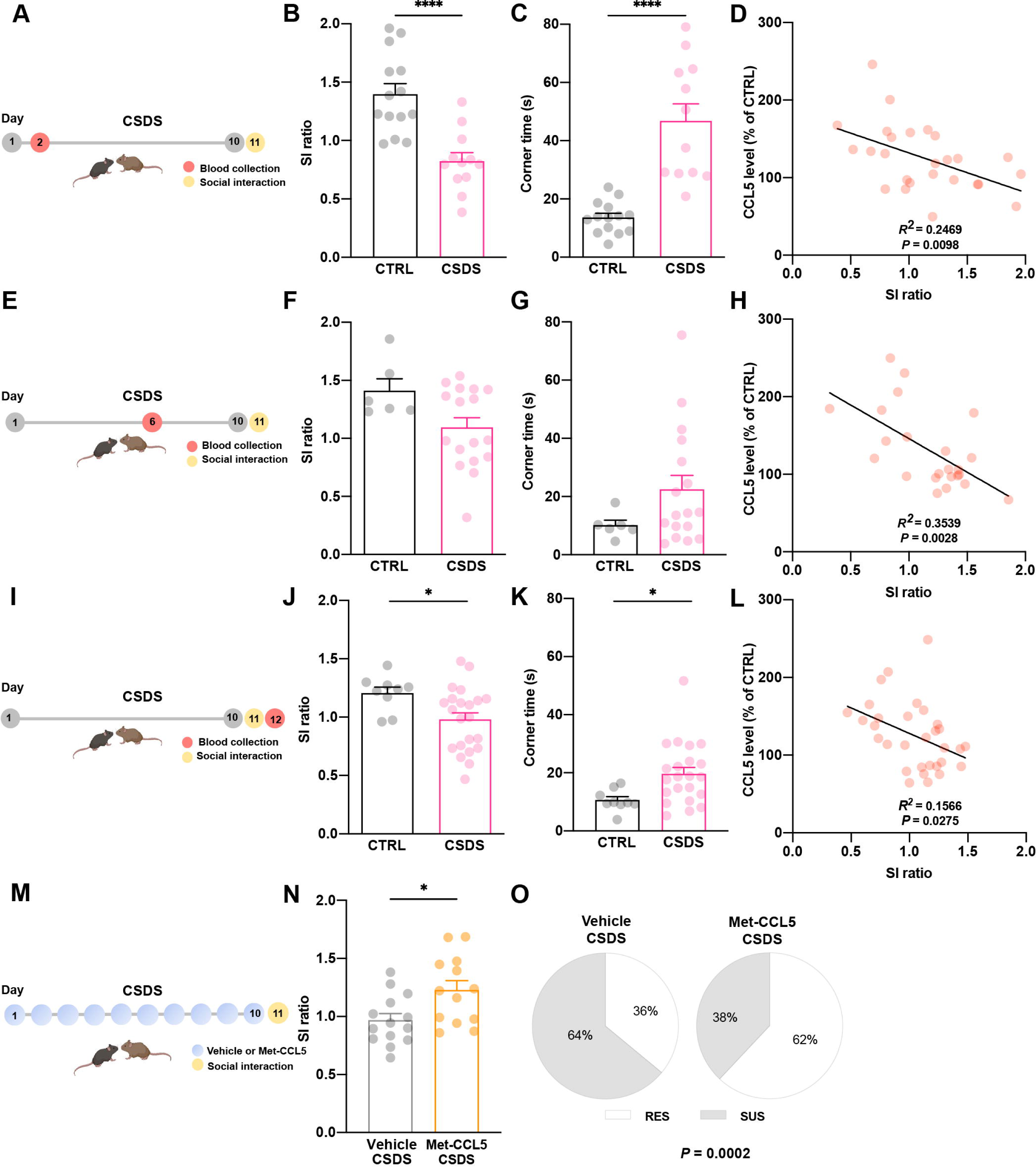
Plasma CCL5 levels correlate with behavioral phenotypes in female mice following CSDS. (A) Experimental design for cohort with blood drawn on day 2. (B) SI ratio from social interaction test, Two-tailed t-test, t(24) = 4.771, *P* < 0.0001; n = 14 (CTRL), 12 (CSDS). (C) Corner zone occupancy duration, Two-tailed t-test, t(24) = 5.905, *P* < 0.0001. (D) Correlation between plasma CCL5 levels and SI ratio, Pearson’s correlation, *R*^2^ = 0.2469, *P* = 0.0098. (E) Experimental design for cohort with blood drawn on day 6. (F) SI ratio, Two-tailed t-test, t(21) = 2.041, *P* = 0.0540; n = 6 (CTRL), 17 (CSDS). (G) Corner zone occupancy duration, Two-tailed t-test, t(21) = 1.474, *P* = 0.1552. (H) Correlation between plasma CCL5 levels and SI ratio, Pearson’s correlation, *R*^2^ = 0.3539, *P* = 0.0028. (I) Experimental design for cohort with blood drawn on day 12. (J) SI ratio, Two-tailed t-test, t(29) = 2.326, *P* = 0.0272; n = 9 (CTRL), 22 (CSDS). (K) Corner zone occupancy duration, Two-tailed t-test, t(29) = 2.495, *P* = 0.0186. (L) Correlation between plasma CCL5 levels and SI ratio, Pearson’s correlation, *R*^2^ = 0.1566, *P* = 0.0275. (M) Experimental design for Met-CCL5 study. Mice were i.p. injected daily with Met-CCL5 or vehicle throughout the CSDS and SI. (N) SI testing in vehicle or Met-CCL5 treated mice following 10-day CSDS, Two-tailed t-test, t(25) = 2.655, *P* = 0.0136. n = 14 (vehicle), 13 (Met-CCL5). (O) Chi-square analysis of percentage of SUS and RES in vehicle and Met-CCL5 treated groups following CSDS, Contingency test, *P* = 0.0002. Data represent mean ± SEM. \**P* < 0.05, \*\**P* < 0.01, \*\*\**P* < 0.001, \*\*\*\**P* < 0.0001.

To assess the longitudinal changes of CCL5 during CSDS and SI testing, we measured the plasma levels of CCL5 in two separate cohorts of female mice on day 6 of the CSDS, which is half way into the CSDS, and 1 day after SI testing (**Fig. 1E-1L**). We found that similar to the CCL5 levels observed one day after the first bout of defeat, SI ratios were negatively correlated with the levels of CCL5 **(Fig. 1H for day 6 and 1L for day 12)**, suggesting that CCL5 may be linked to stress susceptibility following CSDS in female mice. The levels of CCL5 were also significantly correlated with corner zone occupancy on day 6 (**Extended Data** Fig. 1B).

To evaluate the role of peripheral CCL5 signaling in female stress responses, we carried out daily injection of Met-CCL5 (0.4 mg/kg/day i.p.), a functional CCL5 antagonist, throughout the 10-day CSDS and tested for behavioral changes (**Fig. 1M**). We found that Met-CCL5 treatment alone had no effect on SI behavior in unstressed mice. However, daily injection of Met-CCL5 significantly attenuated social avoidance behavior compared to vehicle-injected mice following CSDS (**Fig. 1N**). Chi-square analyses also showed that there was a significant difference in the number of mice exhibiting a resilient phenotype between the two groups (**Fig. 1O**), suggesting that peripheral CCL5 promotes stress-susceptibility in female mice.

## CSDS upregulates microglial CCR5 expression in the PFC in SUS mice

CCL5 has several receptors including CCR1, CCR3 and CCR5. To investigate CCL5 related changes in the brain following CSDS, we conducted RNA-sequencing (RNA-seq) from the nucleus accumbens (NAc) and PFC, the two brain regions known to play important roles in stress and depression. We did not find any CCL5 related changes except for a significant increase of *Ccr5* (∼ 35%) in the PFC in defeated female mice compared to the non-defeated controls (**Fig. 2A** and **Extended Data** Fig. 2A). CCR5 is a seven-transmembrane G protein-coupled receptor expressed in microglia, astrocytes and neurons in diverse brain regions^47–50^. We did not observe similar changes in male mice following CSDS, implicating female specific response of *Ccr5* to CSDS. To assess for clinical relevance, we cross-examined the RNA-seq data of postmortem brain samples from depressed and healthy control human subjects of both sexes published by Labonte *et al*^51^. We found a 2.8-fold (*P* = 0.0078) increase of *Ccr5* expression in the ventromedial PFC (vmPFC) in female MDD subjects compared to the healthy controls (**Fig. 2A**) while no differences were observed in male MDD subjects, suggesting that increased *Ccr5* expression in female mice following CSDS may recapitulate an important and unique feature of depression in women.

**Fig. 2.**
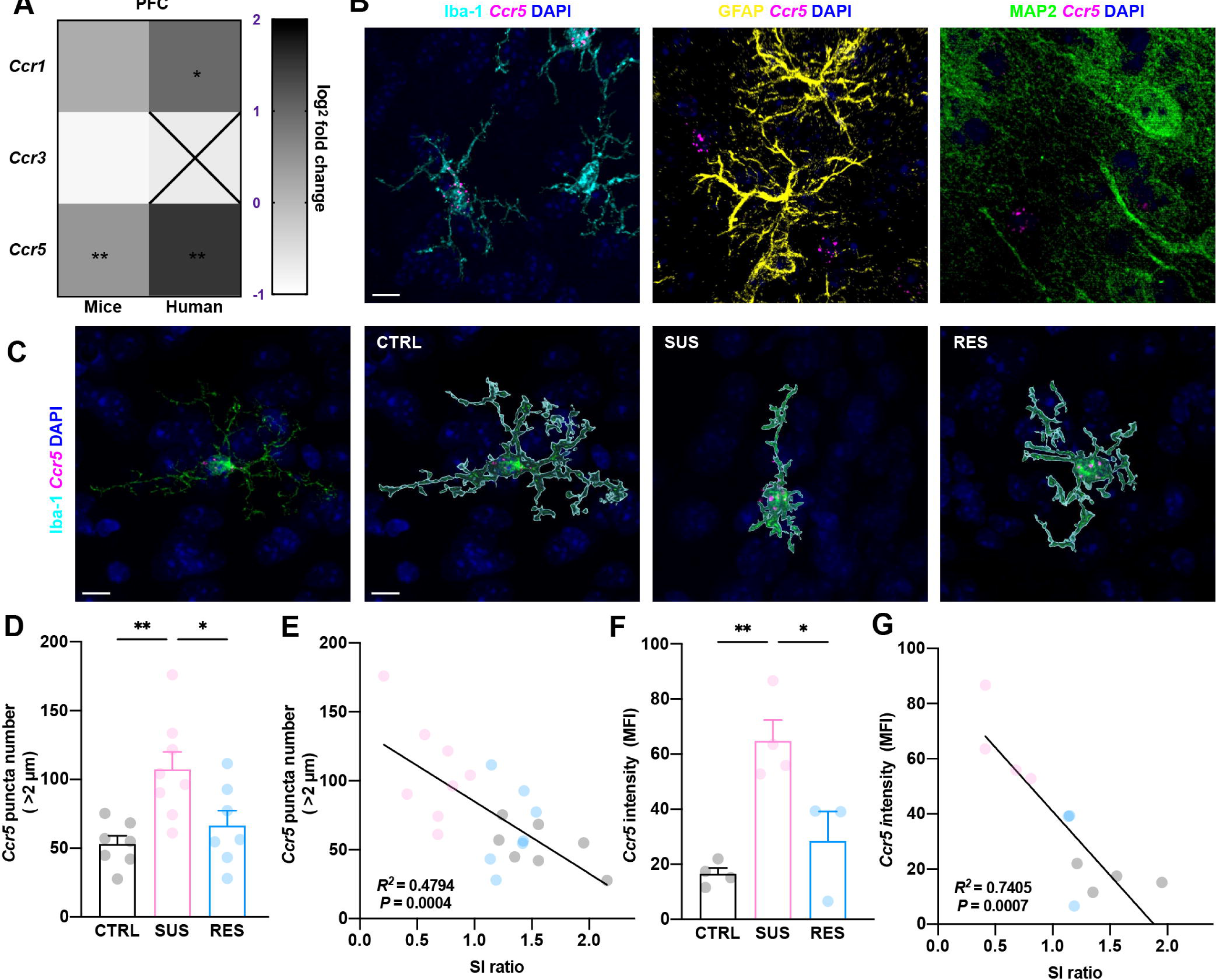
Microglial CCR5 expression in the PFC correlates with stress susceptibility. (A) RNA-seq results of *Ccr1*, *Ccr3* and *Ccr5* expression in the PFC in stressed vs. control female mice and in the vmPFC in female MDD patients vs. healthy controls. Asterisks indicate significance in stressed versus control mice and MDD vs. healthy controls. (B) *Ccr5* expression in the PFC: representative images of *Ccr5* RNA probe with microglia (Iba-1^+^, left), astrocytes (GFAP^+^, middle) and neurons (MAP2^+^, right); scale bar 10 μm. (C) Left: Representative images of *Ccr5* expression in microglia in the PFC of CTRL mice; Scale bar 10 μm. Right: Representative reconstructed images of *Ccr5* puncta in microglia from CTRL, SUS and RES mice. (D) Quantification of *Ccr5* mRNA expression using FISH by calculating *Ccr5*^+^ Iba-1^+^ puncta in CTRL, SUS, and RES mice in the PFC, One-way ANOVA; F(2, 19) = 7.237, *P* = 0.0046; n = 7 (CTRL), 8 (SUS), 7 (RES). (E) Correlation of *Ccr5* puncta with SI ratio, Pearson’s correlation, *R*^2^ = 0.4794, *P* = 0.0004. (F) Quantification of *Ccr5* fluorescence intensity in PFC from CTRL, SUS and RES mice, One-way ANOVA; F(2, 8) = 13.54, *P* = 0.0027; n = 4 (CTRL), 4 (SUS), 3 (RES). (G) Correlation of *Ccr5* fluorescence intensity with SI ratio, Pearson’s correlation, *R*^2^ = 0.7405, *P* = 0.0007. Data represent mean ± SEM. \**P* < 0.05, \*\**P* < 0.01.

To identify the type(s) of cells in the PFC that express *Ccr5,* we performed fluorescent *in situ* hybridization (FISH) combined with immunofluorescence (IF). We only found *Ccr5* mRNA in Iba-1^+^ microglia and did not find any *Ccr5* mRNA signal in either GFAP^+^ astrocytes or MAP2^+^ neurons (**Fig. 2B**). Quantification of *Ccr5* puncta or total *Ccr5* fluorescent signal intensity in the PFC showed significantly increased expression of *Ccr5* in SUS mice compared to the CTRL or RES mice (**Fig. 2C, 2D and 2F**). Correlation analysis confirmed that the number of *Ccr5* puncta (**Fig. 2E**) and total fluorescence intensity of *Ccr5* (**Fig. 2G**) were negatively correlated with SI ratio and positively correlated with time spent in corner zones (**Extended Data** Fig. 2B and 2C). Since CCL5 is the ligand for CCR5 and microglia are the primary cell type in the PFC expressing *Ccr5*, we next tested whether CCL5 can directly influence *Ccr5* expression in microglia-like BV2 cells *in vitro*. We found that incubation of BV2 cells with recombinant CCL5 led to a significant increase in *Ccr5* expression with no effect on *Ccr*1 or *Ccr*3 expression (**Extended Data** Fig. 2D).

## CCR5 inhibition in the PFC attenuates stress-induced neurophysiological and behavioral changes

To investigate the role of CCR5 in the PFC during chronic defeat stress, we inhibited CCR5 expression by introducing lentiviral vectors expressing CCR5-specific shRNA (CCR5-KD) or a scrambled shRNA (Scramble) into the PFC in 6 week-old female mice. Following surgery, mice were given two weeks for recovery and to allow for maximal expression. All mice were subjected to SI testing prior to CSDS (SI-1) to determine the direct effect of viral injection on SI behavior. Mice were then subjected to the 10-day CSDS followed by SI testing (SI-2) and *ex vivo* whole-cell recordings (**Fig. 3A**). CCR5 knockdown was confirmed by quantification of *Ccr5* puncta and qPCR after SI testing (**Fig. 3B** and **Extended Data** Fig. 4A and 4B). We found that CCR5 knockdown in the PFC had no effect on social interaction behavior prior to CSDS (**Fig. 3C**). After 10-day CSDS, CCR5-KD mice had significantly higher SI ratios compared to the mice injected with scrambled shRNA (**Fig. 3D**). Chi-square analyses also showed that more mice in CCR5-KD group exhibited a resilient phenotype compared to the scrambled shRNA injected mice (**Fig. 3E**). Previous studies showed that CCR5 signaling in CA1 neurons decreases neuronal excitability leading to impaired memory processing^52^. Thus, we conducted whole-cell patch-clamp recordings from layer 5 pyramidal neurons in the PFC to assess for the effect of CCR5 knockdown on stress-induced changes in neuronal excitability (**Fig. 3F**). First, we confirmed that CCR5 shRNA injection had no effect on intrinsic neuronal excitability compared to scrambled shRNA injected mice without CSDS. Compared to non-stressed mice, stress induced a significant decrease in the intrinsic membrane excitability in scrambled shRNA injected mice, while knockdown of CCR5 expression significantly prevented this stress-induced decrease seen in L5 PFC pyramidal neurons (**Fig. 3G**). This is also reflected by the measurements of rheobase (**Fig. 3H)**, the minimal injected current needed to elicit an action potential. Following CSDS, pyramidal neurons in scrambled shRNA showed a higher rheobase compared to the other groups (**Fig. 3H**), indicating that the neurons have a decreased intrinsic membrane excitability following CSDS while knockdown CCR5 prevented this stress-induced alteration. Collectively, these data confirm the contribution of CCR5 in the PFC toward stress-mediated behavioral and electrophysiological changes in female mice.

**Fig. 3.**
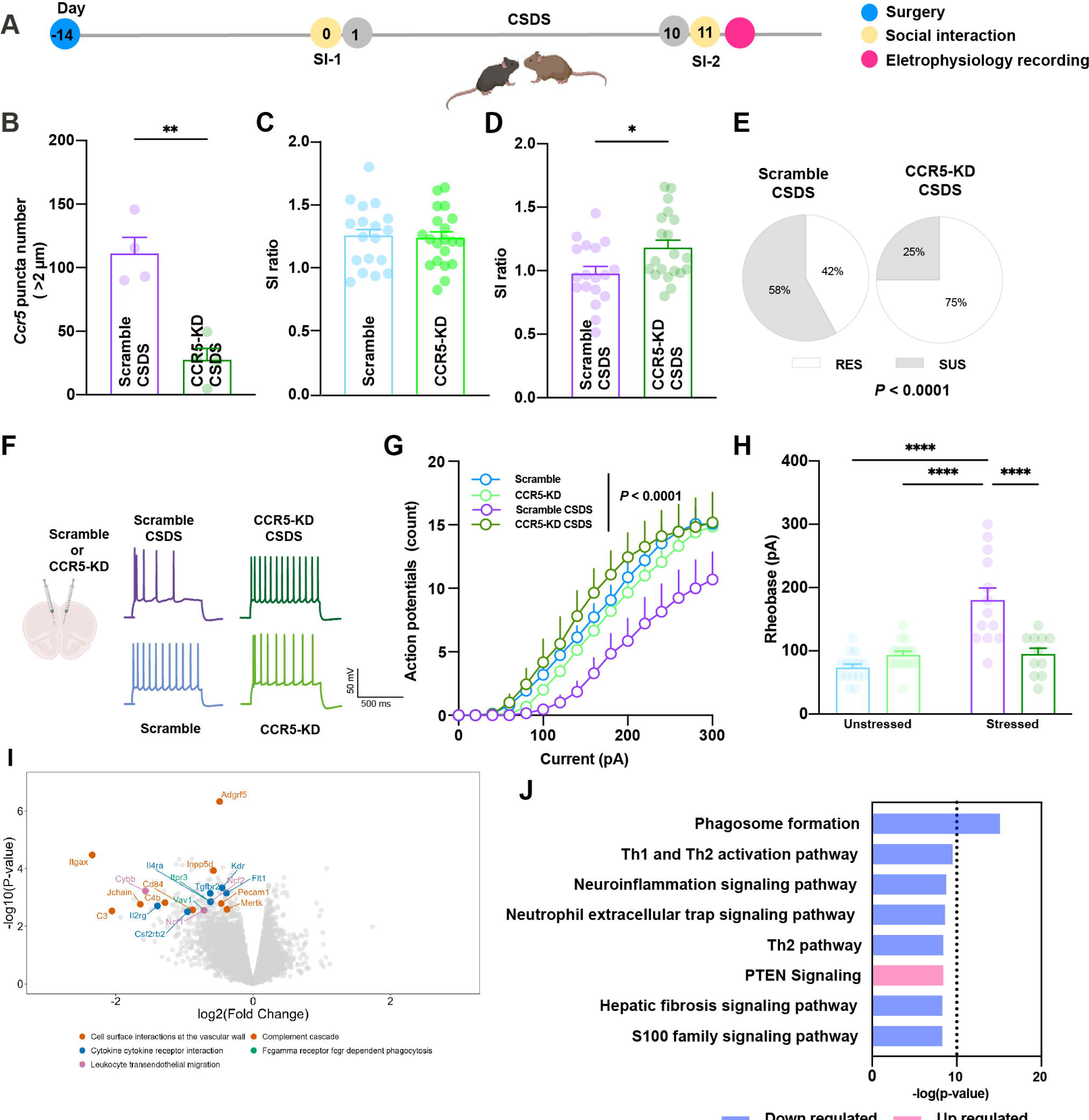
Knockdown CCR5 in the PFC attenuates CSDS-induced social avoidance behavior and intrinsic neuronal excitability alterations. (A) Experimental design. (B) RNA-scope validation of lentivirus–mediated CCR5 knockdown in the PFC, Two-tailed t-test, t(6) = 5.298, *P* = 0.0018, n = 4 (Scramble CSDS), 4 (CCR5-KD CSDS). (C) SI testing in stress naïve females with Scrambled-or CCR5-shRNA injection, Two-tailed t-test, t(36) = 0.2299, *P* = 0.8195; n = 18 (Scramble), 20 (CCR5-KD). (D) SI testing in scrambled-or CCR5-shRNA-injected mice following CSDS, Two-tailed t-test, t(36) = 2.485, *P* = 0.0178; n = 18 (Scramble CSDS), 20 (CCR5-KD CSDS). (E) Chi-square analysis of percentage of RES and SUS mice in scrambled- and CCR5-shRNA-injected mice following CSDS (Contingency test, *P* < 0.0001). (F) Representative traces from *ex vivo* whole-cell current-clamp recordings from scrambled-or CCR5-shRNA injected naïve and stressed mice showing action-potentials generated in response to a 200 pA depolarizing current step. (G) Current-Action potentials curves recorded from PFC L5 pyramidal neurons from scrambled- and CCR5-shRNA injected naïve and stressed mice showing numbers of action potentials elicited by increasing depolarizing current steps, Two way ANOVA, stress x virus manipulation interaction: F(3, 800) = 31.79, *P* < 0.0001, n = 15 (Scramble), 15 (CCR5-KD), 13 (Scramble CSDS), 11 (CCR5-KD CSDS) cells. (H) Rheobase of PFC pyramidal neurons from stressed mice receiving the scrambled-shRNA was significantly higher than the other groups, while CCR5-shRNA blocked this effect in stressed mice, Two-way ANOVA, stress x virus manipulation interaction: F(1,50) = 21.89, *P* < 0.0001, n = 15 (Scramble), 15 (CCR5-KD), 13 (scramble CSDS), 11 (CCR5KD CSDS) cells. (I) Volcano plot showing top 20 DEGs in the PFC in CCR5-KD mice vs. Scrambled mice following CSDS and their GO-term pathway. n = 4 (Scramble CSDS), 6 (CCR5-KD CSDS). (J) Most significantly regulated IPA Canonical Pathways in the PFC from CCR5-KD mice compared to scramble-shRNA-injected mice following CSDS. Data represent mean ± SEM. \**P* < 0.05, \*\**P* < 0.01, \*\*\*\**P* < 0.0001.

To investigate the impact of CCR5 knockdown on stress-induced molecular changes in the PFC, we conducted bulk RNA-sequencing in the PFC. Compared to the scrambled shRNA injected mice, there were 564 downregulated genes and 117 upregulated genes in CCR5-KD mice after CSDS (**Extended Table 2**). Interestingly, the top DEGs are largely involved in cell surface interactions, complement cascade, cytokine and receptor interactions, and leukocyte transendothelial migration (**Fig. 3I)**. Ingenuity pathway analysis (IPA) showed that phagosome formation, Th1 and Th2 activation, and neuroinflammation signaling are the top enriched pathways that are significantly downregulated after CCR5-KD following CSDS **(Fig. 3J)**.

## CSDS induces over-activation of microglia in the PFC in SUS mice

RNA-seq data suggested that CCR5 is involved in regulating immunological processes related to phagosome formation and neuroinflammation during CSDS. Since microglia are the main phagocytic cells in the PFC expressing *Ccr5*, we tested causal roles of microglia CCR5 in these processes. First, we conducted morphological characterization of microglia in the PFC by immunostaining and 3D reconstruction in fixed brain tissue from CTRL, SUS and RES mice. We found that compared to the CTRL or RES mice, microglia in the PFC of SUS mice had shortened process lengths (**Fig. 4A**), and fewer primary branches (**Extended Data** Fig. 3A and 3B), indicating that microglia in SUS mice are more activated compared to those in the CTRL or RES mice. Given that functional analyses identified phagosome formation as the top canonical pathway regulated by CCR5 knockdown. We next examined phagocytic activity in microglia. CD68 is a phagocytic marker and predominantly localizes to lysosomes and endosomes. In resting microglia, CD68 protein is expressed at low levels and its expression increases upon microglial activation^53^. We found that SUS mice had significantly increased amounts of CD68 compared to CTRL or RES mice (**Fig. 4B**). Increased phagocytic activity in activated microglia has been linked to abnormal synaptic pruning under diseased conditions. Thus, we immunolabeled the postsynaptic protein PSD95, an excitatory synaptic marker, and found increased co-localization of PSD95 and CD68 in Iba-1^+^ microglia in SUS mice (**Fig. 4C** and **Extended Data** Fig. 3C), indicating that activated microglia in SUS mice increasingly engulf synaptic material compared to CTRL or RES mice.

**Fig. 4.**
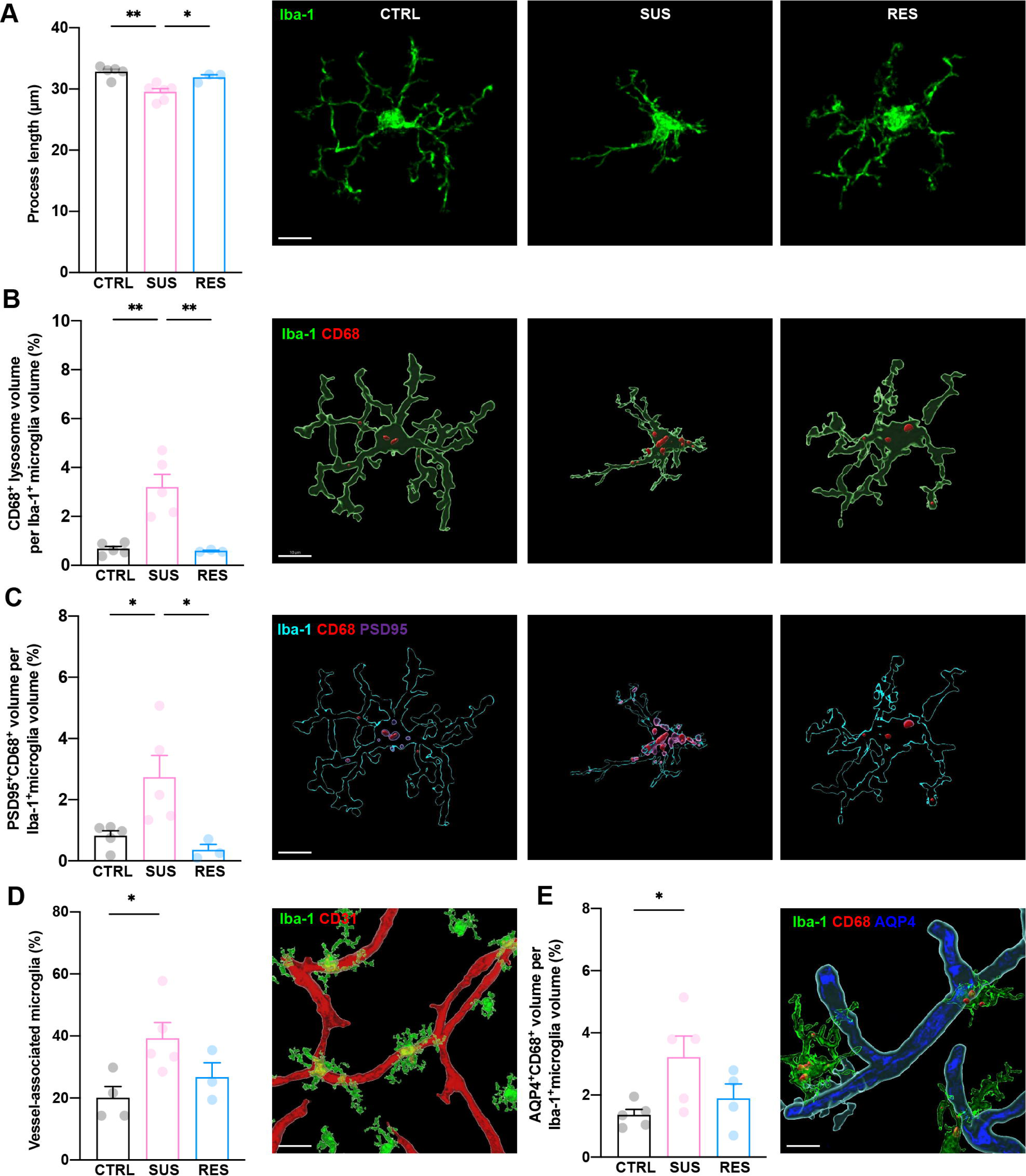
Characterization of microglia in the PFC of CTRL, SUS and RES mice. (A) Quantification of Iba-1^+^ microglia process length in the PFC in CTRL, SUS and RES mice and representative projection of z-stacked images, One-way ANOVA; F(2, 11) = 12.51, *P* = 0.0015, n = 5 (CTRL), 6 (SUS), 3 (RES); scale bar 10 μm. (B) Quantification of CD68^+^ lysosome content inside Iba-1^+^ microglia and representative IMARIS-reconstructed 3D images, One-way ANOVA; F(2, 10) = 16.99, *P* = 0.0006, n = 5 (CTRL), 5 (SUS), 3 (RES); scale bar 10 μm. (C) Quantification of co-localization of synaptic protein PSD95 and CD68 inside Iba-1^+^ microglia in the PFC of CTRL, SUS and RES and representative IMARIS-reconstructed 3D images, One-way ANOVA; F(2, 10) = 6.336, *P* = 0.0167, n = 5 (CTRL), 5 (SUS), 3 (RES); scale bar 10 μm. (D) Quantification of vessel-associated microglia in the PFC of CTRL, SUS and RES, and representative 3D reconstructed image of microglial association with vasculature (CD31: red) in the PFC of SUS mice. One-way ANOVA; F(2, 9) = 4.691, *P* = 0.0402, n = 4 (CTRL), 5 (SUS), 3 (RES); scale bar 20 μm. (E) Quantification of co-localization of AQP4 and CD68 inside Iba-1^+^ microglia in the PFC of CTRL, SUS and RES and representative 3D reconstructed image from microglia (Iba-1: green) engulfing (CD68: red) astrocytic end feet (AQP4: blue) in the PFC of SUS mice. One-way ANOVA; F(2, 11) = 3.991, *P* = 0.0498, n = 5 (CTRL), 5 (SUS), 4 (RES); scale bar 10 μm. Data represent mean ± SEM. \**P* < 0.05, \*\**P* < 0.01.

Functional analyses from RNA-seq data also revealed that CCR5 manipulation influences the expression of genes involved in cell-vascular interaction and leukocyte transendothelial migration, suggesting that microglia maybe associate with the vasculature. To test this, we immunolabeled vasculature with CD31 antibody and microglia with Iba-1, then quantified the number of microglia that are associated with blood vessels. We found that compared to CTRL mice, SUS mice had increased numbers of vessel-associated microglia while RES mice did not significantly differ from the CTRL mice (**Fig. 4D** and **Extended Data** Fig. 3D). To assess whether these vessel-associated microglia increased their phagocytic activity towards components of the neurovascular unit, we immunolabeled the PFC with aquaporin 4 (AQP4)-a transmembrane protein primarily expressed at the end-feet of astrocytes, together with the lysosome marker CD68 and Iba-1, then reconstructed the images using IMARIS software. Quantitative analyses showed that microglia from SUS mice had increased AQP4 inclusion compared to the CTRL mice (**Fig. 4E** and **Extended Data** Fig. 3E), suggesting that vessel-associated microglia indeed participate in the engulfment of vascular components. Interestingly, we did not find significant differences in the amount of AQP4 inclusion between CTRL and RES mice (**Fig. 4E**). Since astrocytic end-feet are integral component of the neurovascular BBB; we then assessed BBB integrity by measuring the expression of claudin 5 (*Cldn5*), a tight junction protein known to control BBB permeability. Consistent with previously published work^54^, we found SUS mice had lower *Cldn5* expression compared to CTRL mice, and there was a positive correlation between *Cldn5* expression and SI ratio (**Extended Data** Fig. 3F) and a negative correlation with corner occupancy duration (**Extended Data** Fig. 3G). These observations suggest that in SUS mice, CSDS induces microglia migration to the cerebral vasculature and increases phagocytosis of vascular materials including astrocytic end-feet that may contribute to the impairment of BBB integrity.

## Down-regulation of CCR5 in the PFC attenuates CSDS-induced microglia activation

To assess the role of CCR5 in CSDS-induced microglial activation in the PFC, we immunostained microglia with Iba-1 and assessed their morphology by 3D reconstruction in CCR5-KD and scrambled shRNA-injected mice following 10-day CSDS. We found that microglia in the PFC of CCR5-KD had longer process lengths and more primary branches compared to those injected with the scrambled shRNA **(Fig. 5A** and **Extended Data** Figs. 4C and 4D). Quantification of the CD68^+^ lysosome content inside microglia in the PFC revealed that CCR5-KD significantly reduced the amount of CD68 per microglia compared to the scrambled shRNA (**Fig. 5B**). Moreover, CCR5-KD mice also reduced the co-localization of PSD95 and CD68 inside Iba-1^+^ microglia (**Fig. 5C** and **Extended Data** Fig. 4E), suggesting attenuated synaptic engulfment by microglia. 3D reconstruction of microglial cell bodies, astrocytic end-feet and vessel structures found that CCR5 knockdown decreased the number of vessel-associated microglia and reduced co-localization of AQP4^+^ with CD68 in vessel-associated microglia **(Figs. 5D and 5E** and **Extended Data** Fig. 4F-G). These studies demonstrate that stress-induced microglial activation and associated engulfment of synaptic material and vascular components are, in part, mediated by CCR5. Functional annotation of RNA-seq data in CCR5-KD PFC confirmed that networks involving abnormal morphology of CNS, phagocytosis and vascular lesions were significantly attenuated in CCR5-KD mice compared to scrambled mice following CSDS (**Fig. 5F** and **Extended Data** Figs. 5 and 6).

**Fig. 5.**
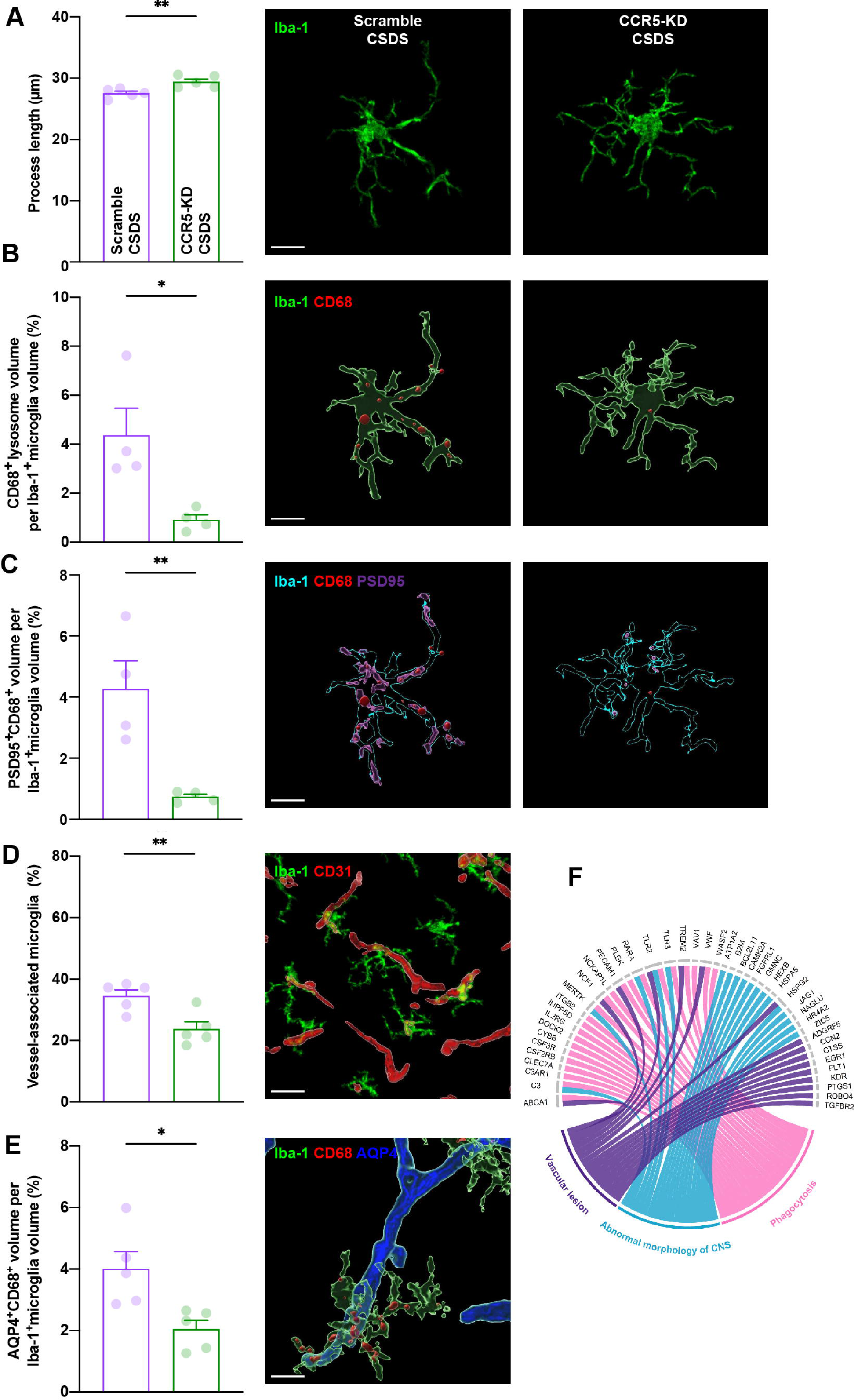
Knockdown CCR5 in the PFC attenuates CSDS-induced microglial over-activation. (A) Quantification of microglial process length in the PFC of scrambled- and CCR5-shRNA injected mice following CSDS and representative z-stacked images. Two-tailed t-test, t(8) = 3.413, *P* = 0.0092, n = 5 (Scramble CSDS), 5 (CCR5-KD CSDS); scale bar 10 μm. (B) Quantification of phagocytosis based on CD68^+^ content in Iba-1^+^ microglia from the PFC of scrambled- and CCR5-shRNA-injected mice following CSDS and representative IMARIS-generated images. Two-tailed t-test, t(6) = 3.091, *P* = 0.0214, n = 4 (Scramble CSDS), 4 (CCR5-KD CSDS); scale bar 10 μm. (C) Quantification of co-localization of synaptic protein PSD95 and CD68 inside Iba-1^+^ microglia in the PFC and representative IMARIS-generated images. Two-tailed t-test, t(6) = 3.843, *P* = 0.0085, n = 4 (Scramble CSDS), 4 (CCR5-KD CSDS); scale bar 10 μm. (D) Quantification of microglia in contact with CD31^+^ blood vessels in the PFC and representative 3D reconstructed image of Scamble-shRNA injected mice following CSDS. Two-tailed t-test, t(8) = 3.417, *P* = 0.0091, n = 5 (Scramble CSDS), 5 (CCR5-KD CSDS); scale bar 20 μm. (E) Quantification of astrocytic end-feet marker AQP4 within CD68^+^ Iba-1^+^ microglia in the PFC and representative 3D reconstructed image of Scamble-shRNA injected mice following CSDS. Two-tailed t-test, t(8) = 3.08, *P* = 0.0151, n = 5 (Scramble CSDS), 5 (CCR5-KD CSDS); scale bar 10 μm. (F) The Circos plot of genes involved in the functional networks of abnormal morphology of CNS, phagocytosis and vascular lesion. Data represent mean ± SEM. \**P* < 0.05, \*\**P* < 0.01.

## Discussion

In current study, we observed that female mice exhibit heightened peripheral inflammatory responses following CSDS. Specifically, we identified CCL5, a C-C motif chemokine, to be persistently elevated in a subset of female mice throughout the duration of CSDS and SI testing. Compared to male mice, where plasma levels of IL-6 predict individual stress susceptibility^46^, plasma levels of CCL5 measured at various time points in females were consistently correlated with stress-susceptibility and blocking CCL5 signaling in the periphery led to increased resilience against CSDS.

In the brain, we found increased expression of the CCL5 receptor CCR5, specifically in microglia in the PFC of SUS mice. Upregulation of CCR5 was associated with: 1) activation of microglia, 2) increased phagocytosis of synaptic material by microglia, 3) increased migration of microglia to the blood vessels, and 4) increased phagocytosis of vascular components. These changes were also associated with neurophysiological changes and impairment of BBB integrity in the PFC. Inhibiting CCR5 signaling in the PFC did not alter intrinsic neuronal excitability or affective behavior in the absence of stress. However, in the context of CSDS, blocking CCR5 signaling directly in PFC attenuated CSDS-induced microglial activation and phagocytic activity, while increasing social behavior and rescue CSDS-induced neuronal hypoexcitability. Together these data show an important role for PFC CCR5 signaling in stress susceptibility in female mice. While synaptic elimination is a critical process for normal brain development and plasticity, aberrant synaptic pruning has been implicated in the pathogenesis of various neuropsychiatric disorders. Reduced numbers of dendritic spines and disrupted network connectivity in various brain regions, including PFC, have been reported both in human patients with MDD and PTSD^55^, as well as in animal stress models^56–58^. Our studies provide evidence that in females, one potential mechanism underlying stress-induced changes in neuronal activity is due to excessive phagocytosis of synaptic materials by overactive microglia in the PFC. Moreover, we demonstrate that this process is largely mediated by microglia specific CCR5 signaling, and that knockdown CCR5 in the PFC significantly reduces microglial phagocytosis of synapses, prevents neuronal excitability changes, and rescues social avoidance behavior in female mice. CCR5 has been implicated in learning and memory through modulation of signaling transduction and neuronal plasticity^59^. In the CA1 region, CCR5 negatively regulates CREB activation and neuronal excitability^52,60^. Our finding that stress-induced increase of CCR5 in the PFC led to neuronal hypoexcitability is consistent with the proposed role of CCR5 in modulating neuronal activity. Previous studies have demonstrated that CX3CL1-CX3CR1 signaling^61^ and the C1q-initiated classical complement cascade^62^ are key molecular pathways involved in the recognition and phagocytosis of synapses by microglia. More recently, TREM2 and its upstream signaling molecule CD33 have been linked to synaptic loss in Alzheimer’s disease^63,64^. RNA-seq data from our study showed that in the PFC, CCR5 knockdown reduces expression of the microglia-specific CX3CR1 as well as molecules involved in the complement pathway, including C1q, C3, and CR3A. Both CD33 and TREM2 were also significantly downregulated compared to scrambled shRNA-injected control mice, suggesting that microglia-specific CCR5 in the PFC may participate in regulating pathways related to excessive synaptic elimination in stress-susceptible mice.

Neurovascular dysfunction and BBB hyperpermeability are linked to psychiatric disorders^65^. Chronic stress-induced downregulation of tight junction proteins and infiltration of peripheral immune components are associated with increased social avoidance behavior in male and female mice^66,67^. It has been proposed that oxidative stress and neuroinflammation contribute to vascular dysfunction and increased BBB permeability^68,69^. Activated microglia produce and secrete proinflammatory cytokines and chemokines, such as TNF-α, IL-1β, IL-6, and MCP-1, that can activate and damage endothelial cells and astrocytes, leading to the disruption of BBB structure and function^70,71^. In our study, stress-susceptible female mice showed increased microglia migration to blood vessels along with increased microglial engulfment of AQP4, a crucial protein enriched in astrocytic end-feet that form the secondary layer of the BBB. This result is consistent with the clinical observation that the coverage of blood vessels by AQP4 is significantly reduced in the mPFC in human patients with MDD^72^. We also demonstrated that CCR5 plays an essential role in this process, and that CCR5 knockdown in the PFC significantly prevented microglia migration and excessive engulfment of AQP4. The current study is the first to demonstrate that activated microglia may interfere with vasculature integrity by phagocytosing neurovascular unit under chronic stress. Moreover, our RNA-seq data suggests that downregulation of CCR5 in microglia may reduce mononuclear cell movement, immune cell adhesion, chemotaxis, and expression of adhesion molecules, including CD11b/CD18 and vascular cell adhesion molecule-1 (VCAM-1), all of which could contribute to reduced microglia migration to blood vessels in CCR5 knockdown mice following CSDS. The exact mechanisms that regulate microglia recruitment to blood vessels and interaction with the vasculature and other immune cells under chronic stress remain to be elucidated.

Previous studies have reported that microglia are selectively activated in certain brain regions, and that resulted in a positively correlation between morphological changes in microglia and their activation status^39,73^. Moreover, causal roles of microglia in stress-mediated affective behavioral changes in previous studies were demonstrated by whole-brain microglia depletion through oral administration of either a CSF1R antagonist or minocycline^74,75^. Because microglia play such an important role in maintaining brain homeostasis by actively surveying the surrounding environment, regulating synaptic activity, results from whole-brain microglia depletion may be difficult to interpret. Our study modulates microglia activity specifically in the PFC while sparing other brain regions that might have opposing effects on stress susceptibility. Moreover, we identified CCR5 as a key molecule mediating microglia activation following CSDS where knockdown of CCR5 effectively modulated multiple pathways and limits stress-induced over-activation of immune responses in the brain. Thus reducing or preventing microglia hyper-activation in the PFC could play an important role in control maladaptive stress responses.

In our study, elevated peripheral CCL5 and PFC CCR5 following CSDS were only found in female, but not male mice, implicating sex-dimorphic responses to stress. In humans, higher plasma levels of CCL5 were found in subjects with PTSD and MDD, and higher concentrations were reported in women compared to men^76,77^. A similar observation was reported in subjects with generalized anxiety disorder and personality disorders^78^. Recent transcriptional analyses also showed increased expression of CCR5 in the PFC of female, but not male, MDD subjects^51^. Changes in CCL5/CCR5 signaling were also associated with treatment efficacy in stress-related disorders in human, with antidepressant treatment response associated with significant reductions of CCL5 and CCR5^79,80^. Furthermore, CCR5 antagonist maraviroc or cenicriviroc treatment also led to improvements in neuropsychological test performance in HIV patients^81–83^. Collectively, these human studies corroborate our studies in mice, supporting the roles of CCL5 and CCR5 in the pathophysiology of stress disorders in females.

In summary, our study investigated the molecular mechanisms underlying stress-induced peripheral and central inflammation. We highlighted a sex-specific role for CCL5 and CCR5 in regulating stress susceptibility in female mice. Our findings provide experimental evidence that targeting the CCL5-CCR5 axis could be a promising therapeutic approach for treating stress-related disorders by preventing aberrant synaptic pruning and preserving neurovascular integrity.

## Materials and Methods

### Animals

Female C57BL/6J mice (6-8 weeks old) and homozygous ERα-Cre male mice on a C57BL/6J background were purchased from the Jackson Laboratory. Female CD-1 mice (Charles River Laboratories) were crossed with male ERα-Cre mice to generate heterozygous ERα-Cre F1 male mice as aggressors. C57BL/6J female mice were housed in groups of 3-5. All mice were maintained on a 12 h light/dark cycle with *ad libitum* access to food and water. Experiments were conducted during the light phase. Procedures were performed in accordance with the National Institutes of Health Guide for Care and approved by the Use of Laboratory Animals and the Icahn School of Medicine at Mount Sinai Institutional Animal Care and Use Committee.

### Female chronic social defeat stress (CSDS)

Female CSDS was performed as previously described^84^. In brief, the F1 male aggressors were injected with CNO (1 mg/kg) 30 min prior to the defeat. Female mice were placed with the aggressor for 5 min of physical defeat and then transferred to their home cage. The procedure was repeated for consecutive 10 days and each day the female mouse encountered a new aggressor.

### Social interaction test

Female mice were placed in a novel interaction open-field arena, with a small animal cage placed at one end. Their movements were monitored and recorded for 2.5 min in the absence (target absent phase), followed by 2.5 min in the presence (target present phase), of a novel non-aggressive F1 male mouse. The total distance traveled, and duration of time spent in the interaction and corner zones were recorded and calculated using EthoVision (Noldus Information Technology Inc). SI behavior was calculated as a ratio of the time spent in the interaction with the target present divided by the time spent in the interaction zone with the target absent. Mice with a ratio above 1.0 were classified as resilient, and mice with a ratio below 1.0 were classified as susceptible.

### Pharmacological treatment with Met-CCL5

Mice were randomly assigned to either vehicle or Met-CCL5 treatment groups. Each mouse received a daily i.p. injections of vehicle (PBS) or Met-CCL5 (0.4 mg/kg) one hour prior to defeat for 10 consecutive days.

### Blood sample collection and CCL5 measurement

Blood samples were collected via the submandibular vein into EDTA-coated tubes (Eppendorf) at different time points. Samples were centrifuged at 2000 *g* for 15 min at 4 °C. Plasma was collected and stored at −80 °C until analysis. Cytokines and CCL5 were measured by Multiplex ELISA (MCYTOMAG-70K, Millipore) or Enzyme Linked Immunosorbent Assay (ELISA, Mouse CCL5/RANTES DuoSet ELISA, #DY478, R&D systems) according to the manufacturer’s instructions.

### Stereotaxic surgery

Six week-old female mice were anesthetized with i.p. injections of ketamine HCl (100 mg/kg) and xylazine (10 mg/kg) and stereotaxically injected with lentiviral vectors carrying either scrambled shRNA or CCR5 shRNA into the PFC (coordinates: AP, +1.8 mm, ML, ±0.75 mm, DV, −2.7 mm; 15° angle). 0.5 μL of virus were bilaterally infused using 33-Gauge Hamilton needles over 5 min, and the needle was left in place for 5 min after the injection.

### Immunofluorescence (IF)

Mice were euthanized by injecting 10% chloral hydrate and transcardially perfused with cold PBS (pH 7.4) followed by 4% paraformaldehyde (PFA). Brains were post-fixed for 12 h in the PFA at 4°C. Coronal sections (40 μm) were prepared on a vibratome (Leica) and subjected to immunofluorescence staining. Specifically, free floating sections were washed with PBS, permeabilized with PBST (PBS + 0.2% Triton X-100) then blocked with 3% normal donkey serum in PBST. Sections were incubated with primary antibodies in 2% normal donkey serum in PBST overnight at 4 °C. Primary antibodies: IBA-1 (1:500, 019-19741, Wako Chemicals), CD68 (1:250, MCA1957, Bio-Rad), MAP2 (1:500, ab5392, Abcam), GFAP (1:500, G3893, Sigma), CD31 (1:300, 102501, Biolegend). Sections were then washed and incubated with secondary antibody (1:400, Jackson ImmunoResearch) for 2 h at RT, then washed 3 times with PBS before staining with DAPI (1 μg/ml, Sigma) for 20 min. Secondary antibodies: Cy2 labeled donkey anti-mouse IgG (H+L) (715-225-150), Cy3-donkey anti-rat IgG (H+L) (712-165-153), Cy5-donkey anti-rabbit IgG (H+L) (711-175-152), all from Jackson ImmunoResearch. Brain sections were then mounted on slides and sealed with coverslips (VWR). Imaging was performed using a Zeiss LSM 780 Confocal Microscope (Zeiss, Oberkochen). 15 μm *z*-stack confocal images were acquired at 1 μm intervals, with 40x/1.3 oil objective or 20x/0.8 at 1x zoom. Image processing was performed using Zen 2011 software (Zeiss). Compressed *z*-stacked immunofluorescence images were used as representative images.

### Morphological analysis of microglia

15-μm *z*-stack images from the PFC on a Zeiss LSM780 Confocal Microscope with 40×/1.3 oil objective at 1× zoom at 1 μm intervals were collapsed into 2D images. Three images were analyzed per animal with *n* = 3-7 mice per condition. ImageJ and Fiji were used to manually trace and generate analyses of process length and number of primary branches of microglia in the PFC. Image quantifications were performed using IMARIS 9.9 software to create surfaces or filaments of each stain based on a threshold applied to all images.

To quantify CD68 and postsynaptic protein PSD95, or AQP4 within microglia, co-localization of CD68 with PSD95 puncta or AQP4 in CD68^+^ phagolysosomes, overlapping surface volume within Iba-1^+^ cell volume was analyzed (expressed as %). To calculate vessel-associated microglia, if the presumed centers of the microglia were overlapping with vessels, the microglia was defined to have contacted the blood vessel (vessel-associated microglia). We then calculated the number of vessel-associated microglia as a percentage of the total microglia in PFC.

### Fluorescent *in situ* hybridization (FISH)

Mice were perfused, brains excised, and fixed in 4% PFA for 24 h at 4°C. Brains were then immersed in 30% sucrose and frozen in optimal cutting temperature embedding media on dry ice. The blocks were sectioned at 15 μm sections and mounted on the slide. FISH was performed using RNAscope Multiplex Fluorescent Detection Reagent Kits v2 (Advanced Cell Diagnostics, #323110) according to manufacturer’s instructions. Briefly, brain sections were first subjected to ISH with nucleotide probe(s) specific for *Ccr5* (RNAscope® Probe - Mm-Ccr5-C3, #438651-C3), followed by standard IF staining with antibodies against astrocytes (GFAP), microglia (Iba-1), or neurons (MAP2). Fluorescent signals were examined with a scanning confocal microscope. *Ccr5* puncta numbers and fluorescent intensity were quantified using a 40× objective lens for each mouse. *Ccr5* in the PFC was imaged in coronal sections by confocal microscopy with a thickness of 10 µm using an SP8 Leica system. The number of puncta was extracted, and density was calculated by dividing the size of region of interest.

### *Ex vivo* electrophysiology

Mice were anesthetized by exposure to isoflurane. Brains were rapidly collected, and coronal sections slices (250 µm thick) were prepared using a Compresstome (VF-300; Precisionary Instruments) in cold (0–4°C) sucrose-based artificial cerebrospinal fluid (SB-aCSF) containing: 87 mM NaCl, 2.5 mM KCl, 1.25 mM NaH_2_PO_4_, 4 mM MgCl_2_, 23 mM NaHCO_3_, 75 mM sucrose, and 25 mM glucose. After 10 min incubation at 32°C for recovery, slices were transferred into oxygenated (95% CO_2_/5% O_2_) aCSF containing: 130 mM NaCl, 2.5 mM KCl, 1.2 mM NaH_2_PO4, 2.4 mM CaCl_2_, 1.2 mM MgCl_2_, 23 mM NaHCO_3_, and 11 mM glucose at room temperature, then transferred to a recording chamber continuously perfused at 2-3 mL/min with oxygenated aCSF. Patch pipettes (4-7 MΩ) were pulled from thin wall borosilicate glass using a micropipette puller (P-97, Sutter Instruments) and filled with a K-Gluconate based intra-pipette solution containing: 116 mM KGlu, 20 mM HEPES, 0.5 mM EGTA, 6 mM KCl, 2 mM NaCl, 4 mM ATP, 0.3 mM GTP and 2 mg/mL biocytin (pH adjusted to 7.2). Cells were visualized using an upright microscope with an IR-DIC lens and illuminated with a white light source (Scientifica). Whole-cell recordings of PFC neurons were performed using a patch-clamp amplifier (Axoclamp 200B, Molecular Devices) connected to a Digidata 1550 LowNoise acquisition system (Molecular Devices). Excitability was measured in current-clamp mode by injecting incremental steps of current (0-300 pA, +20 pA at each step). Signals were low pass filtered (Bessel, 2 kHz) and collected at 10 kHz using the data acquisition software pClamp 11 (Molecular Devices). Electrophysiological recordings were extracted and analyzed using Clampfit (Molecular Devices) and sample traces visualized in R (https://www.r-project.org).

### Bulk RNA sequencing and bioinformatic analysis

PFC samples were collected and processed as described^85^. Bilateral punches were collected from 1 mm coronal slices on ice after rapid decapitation and immediately placed on dry ice and stored at −80 °C until use. RNA was isolated by TRIzol homogenization, chloroform phase separation, and the aqueous RNA layer was processed with an RNeasy Micro Kit (Qiagen) according to the manufacturer’s instructions. RNA integrity was assayed using an RNA Pico chip on a Bioanalyzer 2100 (Agilent, Santa Clara, CA), and only samples with RIN > 9 were considered for subsequent analyses. RNA-seq data analysis and bioinformatics quality control was carried out on the raw sequence data coming from the sequencer using FastQC software. This assesses total sequence, reads flagged as low quality, read length, GC content, per base and per tile quality, per sequence quality score, per base content distribution, per sequence GC content, per base N content, sequence length distribution, sequence duplication levels, overrepresented sequence, adaptor content, and Kmer content. Reads of universal low base quality were discarded and reads with certain low quantity bases were trimmed. Following quality control, RNA reads were mapped to mouse reference genome (mm10) using STAR aligner^86^. Gene-level read counts in every gene model and all exons were derived using STAR. We removed genes or exons that were not expressed in any sample, where we define expression as at least 20 mapped reads in the entire data set. Read counts within samples and between samples were then normalized using the edgeR R-package. The limma model was used to examine the differential expression level (log-transformed) between groups^87^.

### Differential expression analyses

Differentially expressed genes were identified using edgeR with a p-value cutoff of 0.05 and logarithm of fold change (logFC) cutoff of 1.2.

### Pathway Enrichment and Gene Ontology (GO) analysis

GO analyses were performed on differentially expressed genes (DEGs) from different contrasts by using Fisher’s Exact Test (FET) with Benjamini-Hochberg (BH) correction. Gene ontology terms were determined using the Database for annotation. The enriched GO terms with BH-corrected FET *P* < 0.05 were considered statistically significant. Additionally, we performed canonical pathway and functional analysis on the differentially expressed genes using QIAGEN IPA (QIAGEN Inc., https://digitalinsights.qiagen.com/IPA).

### Statistical analysis

All statistical tests comparing control vs. test group(s) were done by two-tailed Student’s t-test, one-way ANOVA analysis was followed by Bonferroni post hoc tests for multiple comparisons, and two-way ANOVA analysis was followed by Tukey post hoc tests for multiple comparisons. In all studies, outliers were identified and excluded, and the null hypothesis was rejected at the 0.05 level. Chi-square tests were used to detect differences in the resilience rates between groups. All statistical analyses were performed using Prism 9 (GraphPad Software, San Diego CA).

## Acknowledgments

Funding was provided by I01 BX005722 from the Veteran’s Administration to J.W and RO1 MH104559 from the National Institute of Mental Health (NIMH) to S.J.R. Funding was provided by Canadian Institutes of Health Research (201811MFE-414896-231226) and a NARSAD Young Investigator Award from the Brain and Behavior Research Foundation (30894) to KLC. In addition, J.W. holds positions in the Research and Development unit of the Basic and Biomedical Research and Training Program at the James J. Peters Veterans Affairs Medical Center. We acknowledge that the contents of this manuscript are solely the responsibility of the authors and do not necessarily represent the views of the NIH or the U.S. Department of Veterans Affairs or the United States Government.

## Data availability statement

Data that support the findings of this study are available from the corresponding author upon reasonable request.

## Author contributions

All authors had full access to all the data in the study and take responsibility for the integrity of the data and the accuracy of the data analysis. *Conceptualization*, S.J.R. and J.W.; *Methodology*, HY L., R.D.C., S.J.R. and J.W.; *Investigation*, HY.L., F.C., L.L., R.D.C., C.G., S.G., Y.S., L.F.P., CZ.Y., K.L.C., F.C., J.W., C.M., J.W.; *Formal Analysis*, HY.L., F.C., R.D.C., Q.W., C.S.B., and J.W.; *Resources*: G.W.H., B.Z., S.J.R. and J.W.; *Writing - Original Draft*, HY.L and J.W.; *Writing - Review & Editing*, HY.L., F.C., R.D.C., K.L.C., L.L., L.F.P., G.W.H.; S.J.R.; *Supervision*, G.W.H., B.Z., S.J.R. and J.W.; *Funding Acquisition*, S.J.R. and J.W.

---

**Extended Table 1. Comparison of select inflammatory cytokines between female and male mice one day after the first bout of defeat.** Data represent mean ± STDEV, n=14-15 per group.

**Extended Table 2. Differential gene expression in the PFC from scrambled-or CCR5-shRNA-injected mice following CSDS.** Summary of DEGs in the PFC from CCR5-or scrambled-shRNA-injected female mice after CSDS. Differential gene expression of next-generation RNA sequencing (RNA-seq) data was obtained using a threshold of 20% difference with *P* < 0.05.

**Extended Data Fig. 1.**
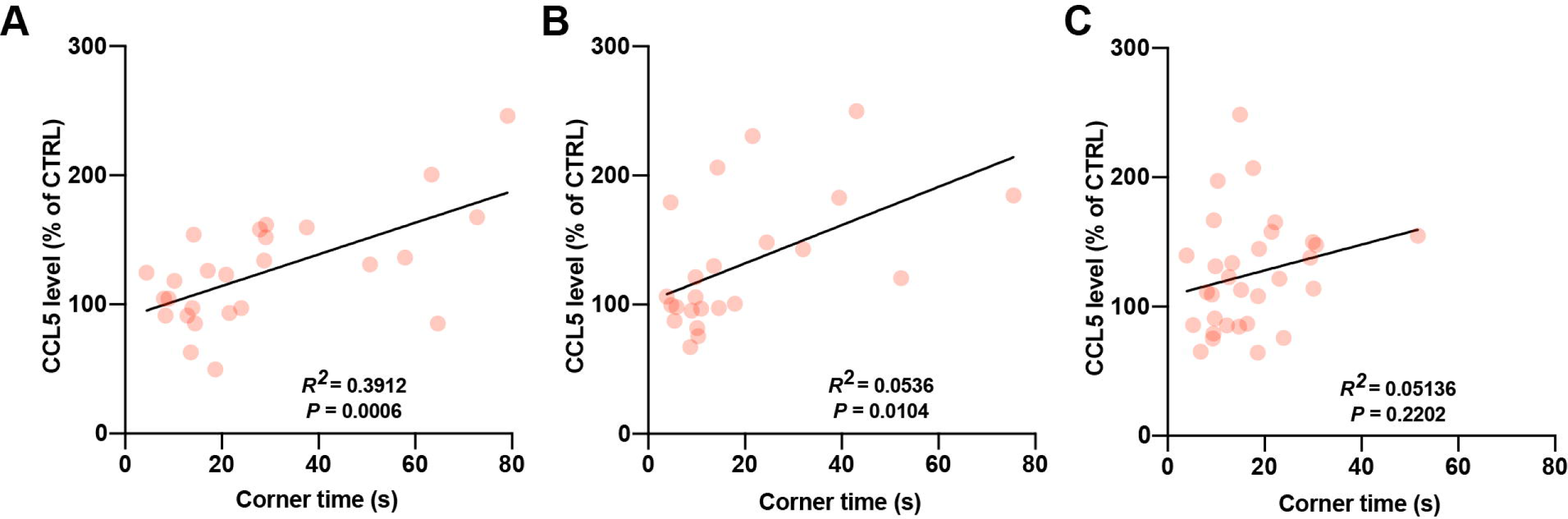
Correlation of plasma CCL5 levels with corner zone occupancy. (A) Correlation of plasma CCL5 levels measured on day 2 and corner zone occupancy duration, Pearson’s correlation, *R*^2^ = 0.3912, *P* = 0.0006. (B) Correlation of plasma CCL5 levels measured on day 6 and corner zone occupancy duration, Pearson’s correlation, *R*^2^ = 0.2738, *P* = 0.0104. (C) Correlation of plasma CCL5 levels on day 12 and corner zone time occupation, Pearson’s correlation, *R*^2^ = 0.05136, *P* = 0.2202. Data represent mean ± SEM.

**Extended Data Fig. 2.**
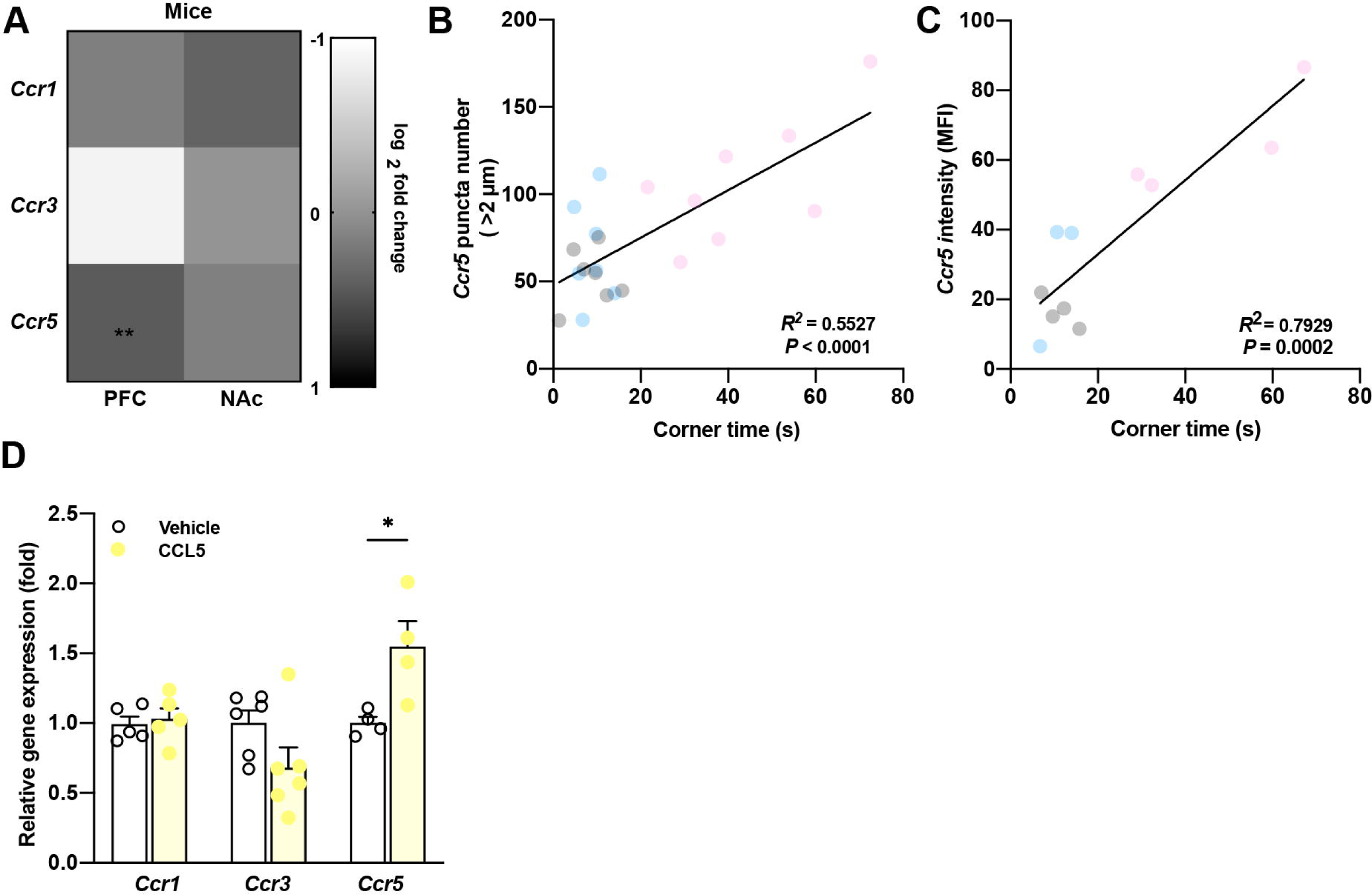
Stress-induced increase of CCR5 expression in PFC is associated with stress-susceptibility. (A) Expression of CCL5 receptors *Ccr1, Ccr3* and *Ccr5* in the PFC and NAc in female mice following CSDS compared to the non-stressed CTRL mice. (B) Correlation of the number of *Ccr5* puncta and corner zone occupancy, Pearson’s correlation, *R*^2^ =0.5527, *P* < 0.0001. (C) Correlation of *Ccr5* fluorescence intensity and corner zone occupation, Pearson’s correlation, *R*^2^ =0.7929, *P* = 0.0002. (D) *In vitro* testing of *Ccr1, Ccr3 and Ccr5* expression in BV2 cells upon stimulation with 20 ng/mL CCL5, Two-tailed t-test, *Ccr1*: t(8) = 0.4009, *P* = 0.699, n = 5; *Ccr3*: t(10) = 1.865, *P* = 0.0917, n = 6; *Ccr5*: t(6) = 2.889, *P* = 0.0277, n = 4. Data represent mean ± SEM. \**P* < 0.05, \*\**P* < 0.01.

**Extended Data Fig. 3.**
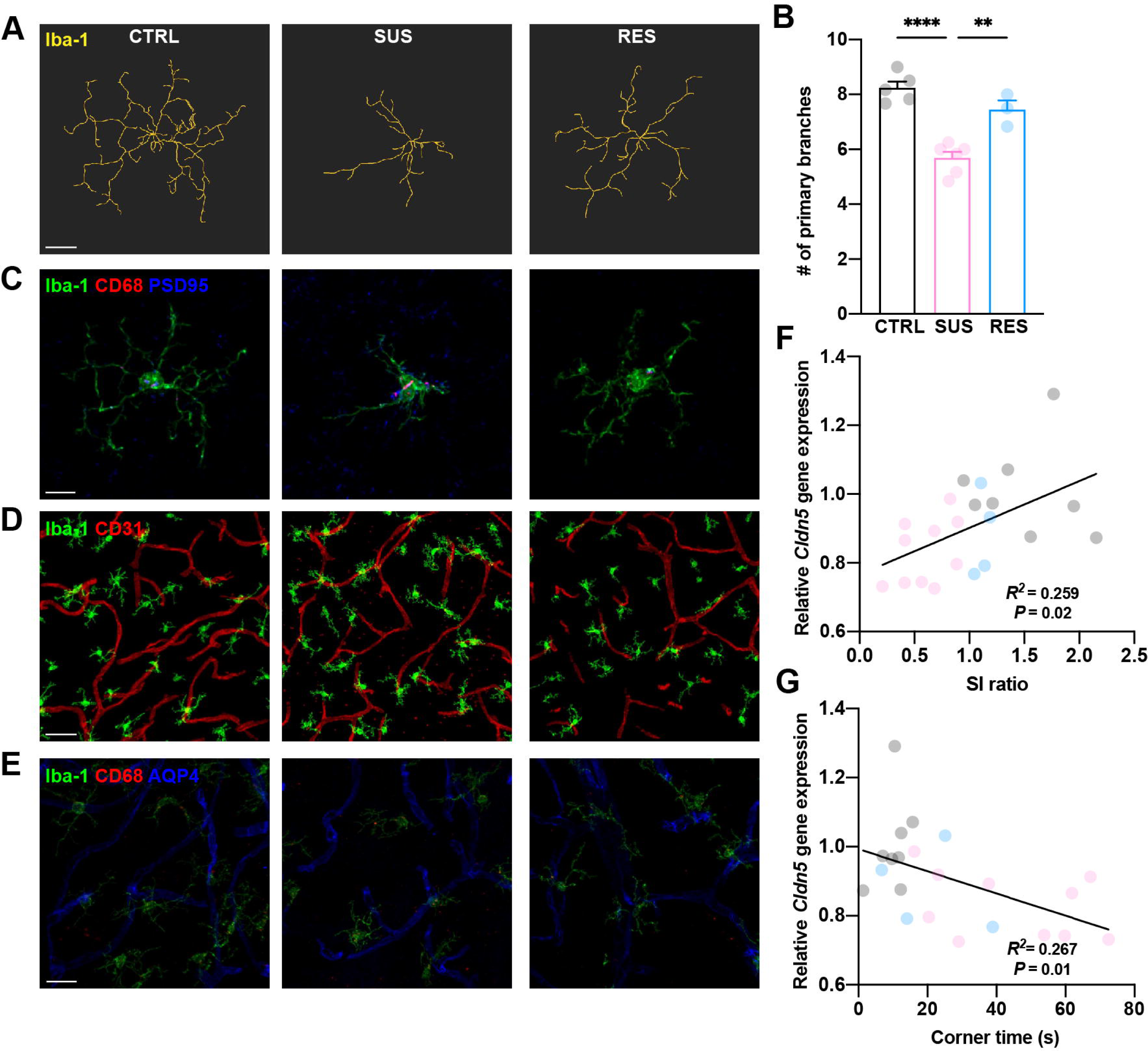
Characterization of microglia in the PFC of CTRL, SUS and RES mice. (A) Representative reconstructed images of microglia surface using IMARIS filament tracer module. (B) Quantification of the number of primary branches in microglia in the PFC of CTRL, SUS and RES mice. One-Way ANOVA, F(2, 11) = 30.47, *P* < 0.0001; scale bar 10 μm. (C) Representative fluorescent images of Iba-1^+^ microglia (green) and CD68 (red) and PSD95 (blue); scale bar 10 μm. (D) Representative fluorescent images of Iba-1^+^ microglia (green) and CD31 (red); scale bar 40 μm. (E) Representative fluorescent images of Iba-1^+^ microglia (green), CD31 (red) and AQP4 (blue); scale bar 20 μm. (F) Correlation of *Cldn5* mRNA expression in the PFC with SI ratio, Pearson’s correlation, *R*^2^ = 0.2585, *P* = 0.0157, n = 8 (CTRL), 10 (SUS), 4 (RES). (G) Correlation of *Cldn5* expression with corner time occupancy, Pearson’s correlation, *R*^2^ = 0.2667, *P* = 0.0139, n = 10 (CTRL), 8 (SUS), 4 (RES). Data represent mean ± SEM. \*\**P* < 0.01, \*\*\*\**P* < 0.0001.

**Extended Data Fig. 4.**
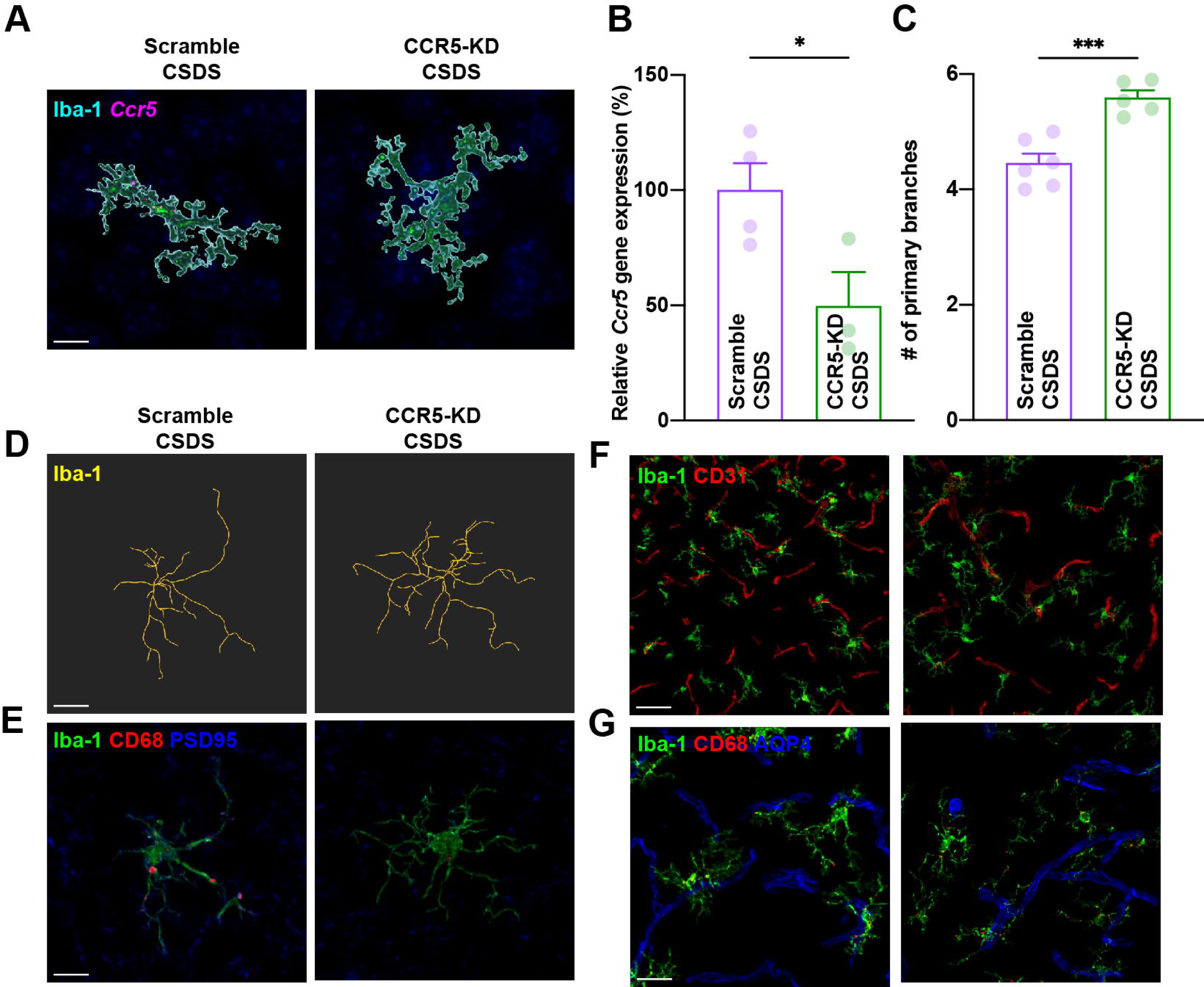
Characterization of microglia in the PFC of scrambled- and CCR5-shRNA-injected mice following CSDS. (A) Representative 3D reconstructed images of PFC microglial *Ccr5* expression in scrambled- and CCR5-shRNA-injected CSDS mice; scale bar 10 μm. (B) qPCR validation of Ccr5 expression in the PFC, Two-tailed t-test, t(5) = 2.704, *P* = 0.0426, n = 4 (scramble CSDS), 3 (CCR5-KD CSDS). (C) Quantification of the number of primary branches in microglia in the PFC of scrambled- and CCR5-shRNA injected mice following CSDS, Two-tailed t-test, t(9) = 5.203, *P* = 0.0006, n = 6 (scramble CSDS), 5 (CCR5-KD CSDS). (D) Representative reconstructed image of microglia surface using IMARIS filament tracer module; scale bar 10 μm. (E) Representative images of PFC immunolabeled with Iba-1 (green), CD68 (red), PSD95 (blue); scale bar 10 μm. (F) Representative images of PFC immunolabeled with Iba-1 (green) and CD31 (red), scale bar 40 μm or (G) Representative images of PFC immunolabeled with Iba-1 (green), CD68 (red) and AQP4 (blue); scale bar 20 μm. Data represent mean ± SEM. \**P* < 0.05, \*\*\**P* < 0.001.

**Extended Data Fig. 5.**
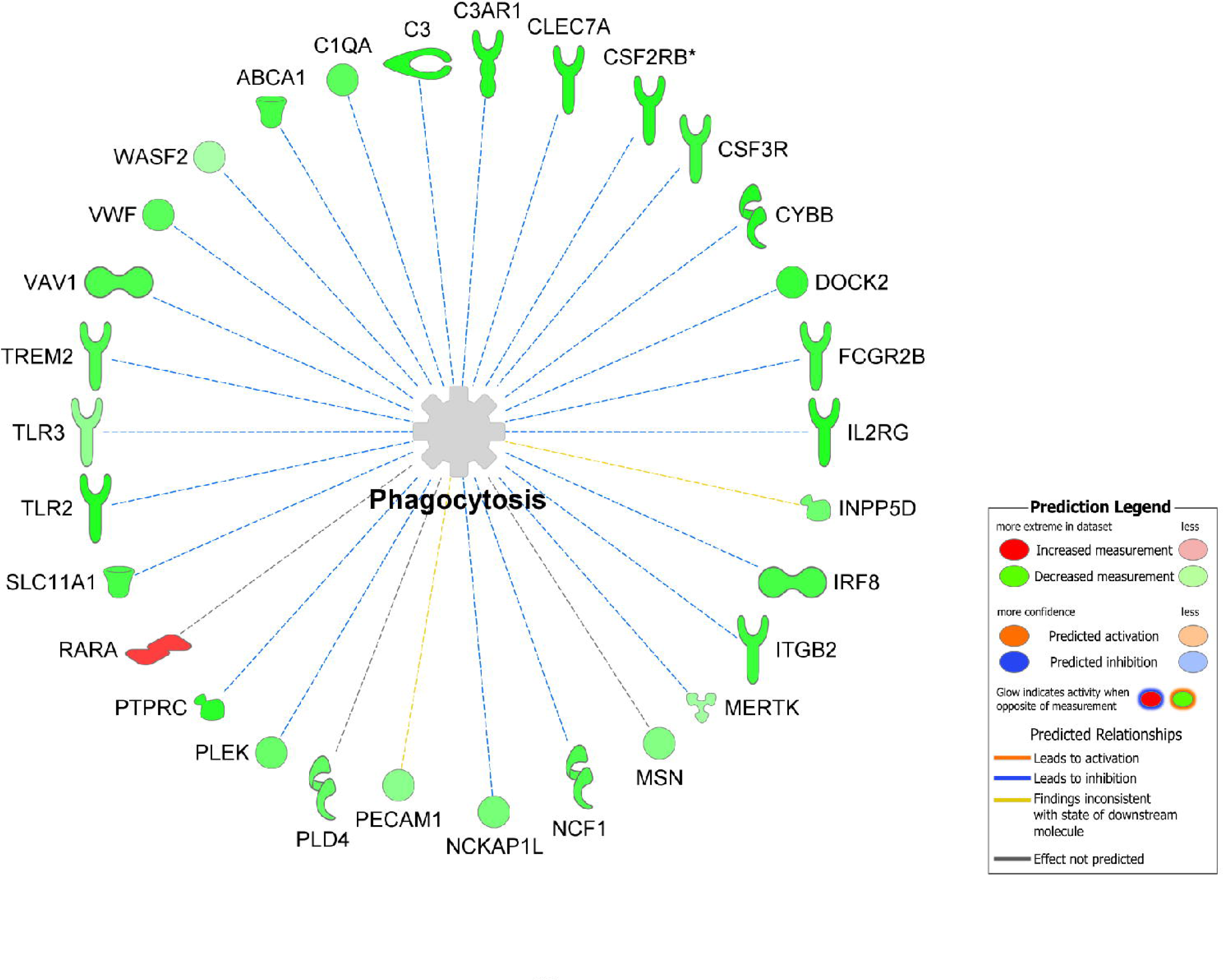
Graphical representation of DEGs annotated with the functional network “phagocytosis” in PFC of defeated female mice. Ingenuity pathway analysis (IPA) was used for prediction of the functional networks of DEGs in the PFC of mice subjected to scramble-or CCR5-shRNA injected mice following CSDS. Among 71 differentially expressed genes involved in the “phagocytosis”, top 30 genes with P < 0.01 were shown. IPA identified the functional network “phagocytosis” to be down-regulated in CCR5-KD CSDS mice. Genes that were downregulated in CCR5-shRNA CSDS relative to scramble CSDS mice are shown in green; genes that were upregulated are shown in red.

**Extended Data Fig. 6.**
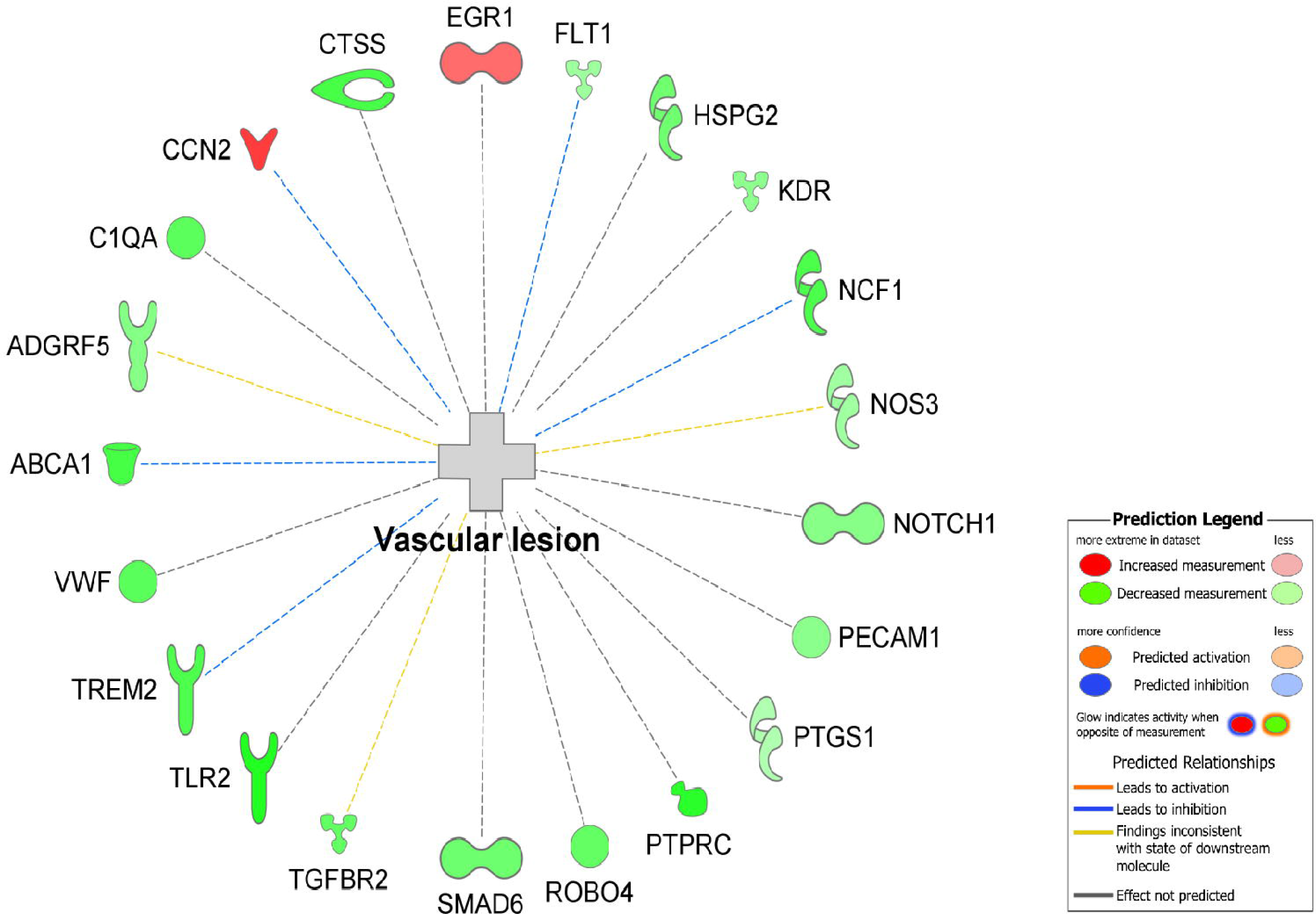
Graphical representation of DEGs annotated with the functional network “vascular lesion” in PFC of defeated female mice. Ingenuity pathway analysis (IPA) was used for prediction of the functional networks of differentially expressed genes in the PFC of mice subjected to scramble-or CCR5-shRNA injected mice following CSDS. Among 51 DEGs involved in the “vascular lesion”, top 21 genes with P < 0.01 were shown. IPA identified the functional network “vascular lesion” to be down-regulated in CCR5-KD CSDS mice. Genes that were downregulated in CCR5-KD CSDS relative to scramble CSDS mice are shown in green; genes that were upregulated are shown in red.

