## Extended Table 1 for "Chemokine receptor 5 signaling in PFC mediates stress susceptibility in female mice"

### Table 1. Characteristic of inflammatory cytokines in female and male mice after first bout of defeat

| | Female | | | Male | | |
|---|---|---|---|---|---|---|
| | CTRL | CSDS | P Value | CTRL | CSDS | P Value |
| CCL5 | 16.6 (5.8) | 24.7 (7.2) | 0.0026 | 18.4 (7.5) | 18.8 (6.2) | >0.05 |
| IL-1α | 100.4 (50.0) | 257.4 (151.8 ) | 0.0008 | 124.2 (29.9) | 195.0 (143.7) | >0.05 |
| IL-1β | 2.9 (1.1) | 7.2 (6.15) | 0.013 | 6.9 (6.2) | 13.6 (16.8) | >0.05 |
| IL-6 | 3.2 (2.0) | 67.4 (102.1) | 0.0266 | 2.7 (3.5) | 34.4 (32.0) | 0.0201 |
| IL-12 (p70) | 7.8 (6.2.2) | 36.5 (6.2) | 0.0002 | 7.7 (6.6) | 15.7 (22.6) | >0.05 |
| MCP-1 | 16.7 (17.7) | 162.9 (181.9) | 0.0045 | 13.5 (6.0) | 25.2 (20.1) | >0.05 |
| TNF-α | 4.3 (1.3) | 15.4 (11.0) | 0.0006 | 1.7 (1.3) | 3.9 (4.3) | >0.05 |
