## Extended Table 2 for "Chemokine receptor 5 signaling in PFC mediates stress susceptibility in female mice"

**Upregulated genes in CCR5-KD CSDS versus scramble-CSDS mice**

| Gene | logFC | PValue | FDR |
|---|---|---|---|
| Hikeshi | 0.26341039 | 0.00997719 | 0.38444445 |
| Cox7c | 0.26344003 | 0.01308369 | 0.40977394 |
| Nkrf | 0.26459497 | 0.00590254 | 0.33152234 |
| Rpf2 | 0.26557173 | 0.02613282 | 0.47277542 |
| Eif1a | 0.26575955 | 0.01238603 | 0.40296079 |
| Zfp14 | 0.26697838 | 0.0166038 | 0.41913632 |
| Cox6c | 0.26730375 | 0.02080777 | 0.45620354 |
| Mtx2 | 0.26749076 | 0.00133485 | 0.25500345 |
| Acyp2 | 0.26769084 | 0.03660155 | 0.51547555 |
| Hibch | 0.26863339 | 0.0192937 | 0.44287137 |
| Ndn | 0.26890893 | 0.02042502 | 0.45361522 |
| Ndufa12 | 0.26959427 | 0.00946745 | 0.38444445 |
| Arhgap15 | 0.26965562 | 0.04161725 | 0.53390655 |
| Galnt9 | 0.27056792 | 0.00685666 | 0.36618602 |
| Bud31 | 0.27070523 | 0.00280778 | 0.26893792 |
| Zfp101 | 0.27218014 | 0.03024483 | 0.48937787 |
| Fastkd5 | 0.27357601 | 0.03812302 | 0.52080674 |
| Tomm7 | 0.27373683 | 0.04742722 | 0.54737976 |
| Ndufb2 | 0.27433912 | 0.02220337 | 0.45620354 |
| Tmem97 | 0.27462845 | 0.04756237 | 0.54740929 |
| Hat1 | 0.27582376 | 0.00531572 | 0.32054063 |
| Mterf2 | 0.27617609 | 0.01310544 | 0.40977394 |
| Eef1akmt1 | 0.27805986 | 0.00890706 | 0.38444445 |
| Zfp958 | 0.27829808 | 0.013153 | 0.40977394 |
| Taf7 | 0.27861076 | 0.02350224 | 0.45620354 |
| Mkks | 0.28008883 | 0.00551473 | 0.32054063 |
| Dpy30 | 0.28103235 | 0.00964955 | 0.38444445 |
| Ankrd29 | 0.28615035 | 0.04539948 | 0.53953037 |
| Mrps21 | 0.28663423 | 0.0023359 | 0.26893792 |
| Dusp5 | 0.28724525 | 0.02269735 | 0.45620354 |
| Banp | 0.28826034 | 0.03502678 | 0.51103087 |
| Uqcrh | 0.28926797 | 0.0029119 | 0.26893792 |
| Pou3f1 | 0.29230904 | 0.03511117 | 0.5117462 |
| Fezf2 | 0.29361699 | 0.01674526 | 0.41913632 |
| Tomm5 | 0.29366783 | 0.00460345 | 0.30955422 |
| Fam162a | 0.2951531 | 0.02152754 | 0.45620354 |
| Lratd2 | 0.29624701 | 0.04093319 | 0.5303314 |
| Lmo3 | 0.29979733 | 0.00054032 | 0.20874477 |
| Ndufa4 | 0.30106349 | 0.0026835 | 0.26893792 |
| Peg10 | 0.30527515 | 0.00063837 | 0.20874477 |
| Zswim5 | 0.30651129 | 0.03910217 | 0.52303674 |
| Coa6 | 0.30667384 | 0.00758531 | 0.37600986 |
| Tmem177 | 0.30733458 | 0.03156095 | 0.49597524 |
| Immp1l | 0.30901806 | 0.00876238 | 0.38313585 |
| Gpr45 | 0.30983982 | 0.00590982 | 0.33152234 |
| Tfb1m | 0.31756907 | 0.04136533 | 0.53296204 |
| Pigx | 0.31796372 | 0.01516709 | 0.41574477 |
| Glra2 | 0.31824682 | 0.02486151 | 0.46207746 |
| Mrpl53 | 0.32257976 | 0.02571784 | 0.467747 |
| Hspa5 | 0.32279916 | 0.00083471 | 0.22536139 |
| Slc35f3 | 0.32317929 | 0.0374435 | 0.5188215 |
| 1500009L16F | 0.32538514 | 0.03577549 | 0.51389656 |
| St7 | 0.32597293 | 0.00870707 | 0.38313585 |
| Atp5mpl | 0.32651541 | 0.01413205 | 0.411293 |
| Cox7b | 0.33026372 | 0.00811601 | 0.37965616 |
| Cabp7 | 0.33104503 | 0.03540798 | 0.51297273 |
| B9d2 | 0.34003424 | 0.01332093 | 0.41045376 |
| Egr1 | 0.34011662 | 0.00055504 | 0.20874477 |
| Msrb1 | 0.34193147 | 0.00484966 | 0.31430344 |
| Olfm3 | 0.34222793 | 0.01374879 | 0.41045376 |
| Arxes1 | 0.34288752 | 0.00094601 | 0.23206128 |
| Uchl3 | 0.34356796 | 0.00037525 | 0.20177257 |
| Cadps2 | 0.35040643 | 0.0357576 | 0.51389656 |
| Galnt14 | 0.35224082 | 0.01166928 | 0.39707572 |
| Arhgap25 | 0.35427165 | 0.04346003 | 0.53592893 |
| Lipo3 | 0.35627038 | 0.02362403 | 0.45620354 |
| Smim17 | 0.35678426 | 0.00206764 | 0.26718678 |
| 3300002P13I | 0.36268469 | 0.00232673 | 0.26893792 |
| Ndufa1 | 0.36702507 | 0.01642641 | 0.41913632 |
| Zfp273 | 0.36775445 | 0.02087188 | 0.45620354 |
| Samd15 | 0.38365043 | 0.03888606 | 0.52256072 |
| Gpr3 | 0.39426735 | 0.01444752 | 0.41164191 |
| Mcm5 | 0.3983606 | 0.0172955 | 0.41999632 |
| Zfp825 | 0.40043316 | 0.01604412 | 0.41864312 |
| Apoo | 0.41076166 | 0.04060791 | 0.5299534 |
| Flywch2 | 0.41408675 | 0.01319715 | 0.40977394 |
| Marchf11 | 0.41486695 | 0.02455452 | 0.45949774 |
| Pdlim1 | 0.41490817 | 0.01526732 | 0.41594516 |
| Pcdhb10 | 0.41862923 | 0.03244946 | 0.50336661 |
| Bcl11b | 0.42946147 | 0.00429761 | 0.30126027 |
| Sstr2 | 0.43067522 | 0.00069969 | 0.20874477 |
| Mis18a | 0.43606096 | 0.01981253 | 0.44734288 |
| Ints12 | 0.4366183 | 0.00039783 | 0.20177257 |
| Rara | 0.44533331 | 0.00379392 | 0.29399113 |
| Lypd6b | 0.44780569 | 0.01894982 | 0.44183226 |
| Ccn2 | 0.44887206 | 0.00020286 | 0.17270148 |
| Plpp4 | 0.44950003 | 0.03268409 | 0.50450416 |
| D230017M19 | 0.45598848 | 0.01468918 | 0.4136119 |
| Skap1 | 0.47373624 | 0.03582114 | 0.51389656 |
| Tmem163 | 0.48110035 | 0.00067541 | 0.20874477 |
| Serinc2 | 0.48161049 | 0.04068205 | 0.52996516 |
| Abracl | 0.49932397 | 0.00087198 | 0.22536139 |
| Tnfrsf23 | 0.51039371 | 0.04256632 | 0.5350035 |
| Pcp4 | 0.51913982 | 0.03957678 | 0.52743533 |
| Ube2cbp | 0.52344282 | 0.02001353 | 0.44873073 |
| Pnldc1 | 0.52590277 | 0.03666838 | 0.51547555 |
| Naaladl2 | 0.53508775 | 0.02888258 | 0.48403804 |
| Fcer2a | 0.56853855 | 0.0454205 | 0.53953037 |
| Gadd45a | 0.5902665 | 0.02381776 | 0.45620354 |
| Klhl1 | 0.59116845 | 0.01936969 | 0.44287137 |
| Slc39a5 | 0.62090835 | 0.0265806 | 0.47611515 |
| Npc1l1 | 0.6453935 | 0.04139082 | 0.53296204 |
| Trh | 0.64884846 | 0.02527818 | 0.46545937 |
| Chrna2 | 0.70038978 | 0.01968156 | 0.44577649 |
| Trhr2 | 0.71506249 | 0.02250898 | 0.45620354 |
| Pappa2 | 0.74596676 | 0.04479354 | 0.53953037 |
| Rslcan18 | 0.75307662 | 0.04213969 | 0.53412906 |
| Gdf9 | 0.76288158 | 0.01386802 | 0.41045376 |
| Kbtbd6 | 0.76558103 | 0.03532831 | 0.51284523 |
| Zfp456 | 0.78103454 | 0.01998375 | 0.44873073 |
| Pkd2l1 | 0.92340187 | 0.00158348 | 0.25500345 |
| Nr4a2 | 0.96755181 | 0.00430131 | 0.30126027 |
| Rxfp2 | 0.99357097 | 0.00281567 | 0.26893792 |
| Acaa1b | 1.01521061 | 0.01657877 | 0.41913632 |
| Gpr139 | 1.01934218 | 0.0420472 | 0.53412906 |
| Dlk1 | 1.07156138 | 9.7558E-05 | 0.13585454 |
| Pkp1 | 1.73590798 | 0.01043153 | 0.38444445 |
